# Emergence of a somatosensory tonotopic map for substrate vibration in the brainstem

**DOI:** 10.1101/2023.09.26.559502

**Authors:** Kuo-Sheng Lee, Dominica de Thomas Wagner, Mark Sanders, Daniel Huber

## Abstract

Perceiving substrate vibrations is a fundamental component of tactile perception. The wide frequency spectrum of vibrations is covered by integrating responses of multiple mechanoreceptors that innervate various subtypes of mechanosensitive end organs, each preferring a specific range: Merkel cells (0.5-10Hz), Meissner corpuscles (10-150Hz) and Pacinian corpuscles (150-1000Hz) in primates. As the density of different end organs greatly varies across the body, each body part potentially has a specific frequency preference. How location (somatotopy) and frequency tuning (tonotopy) are processed along the ascending neuraxis and how they converge to drive responses of individual neurons is poorly understood. In this study, we address this question by combining *in vivo* peripheral electrophysiology and two-photon calcium imaging along the entire dorsal column-medial lemniscal pathway, including the dorsal root ganglia, dorsal column nuclei (DCN), the thalamus and the cortex. Surprisingly, we found that both frequency, as well as location, are organised into structured maps in the DCN. Furthermore, both maps are intimately related at the fine spatial scale with parallel map gradients that are consistent across the depth of the DCN and preserved along the ascending pathway. Additional sensory mapping experiments based on peripheral characterisation revealed that the tonotopic map only partially reflects the distribution of end organs in the skin and deep tissue. Instead, we show that the emergence of the finescale tonotopy is probably due to the selective dendritic sampling from axonal afferents, right at the first synaptic relay. Taken together, we conclude DCN neural circuits are key to the emergence of these two fine-scale topological organisations in early somatosensory pathways. The underlying computational principle is intriguingly similar to the integration of multiple functional maps along the ascending visual pathways, suggesting a universal law governing the optimization of sensory systems.

## Introduction

Vibrations are ecologically relevant sensory stimuli that can convey different types of information such as surface texture, or movement of an object near us ^1,2^. Although substrate vibrations can convey complex information, they are actually composed of only a few physical attributes: frequency, amplitude and the location of stimulation. The ability to feel substrate vibrations, or gentle touch over a broad range of frequencies fundamentally depends on sensory receptors in skin, joints and bones. They are known as low-threshold mechanoreceptors (LTMRs), which innervate various mechanosensitive end organs, and are specialised to respond to different frequencies ^3,4^. For example, Merkel cells and Meissner corpuscles are sensitive to low-frequency vibrations (below 150 Hz), while Pacinian corpuscles are sensitive to higher frequencies (above 150 Hz).

Whereas the different end-organ complexes likely play crucial roles in the mechanotransduction at the site of stimulation, the LTMRs transmit this information to the central nervous system ^5,6^. Most of LTMRs either directly synapse on neurons of the cuneate (upper body) and gracile nuclei (lower body) of the DCN or indirectly connect through spinal cord dorsal horn to the brainstem ^7,8^, from where the information further ascends via the ventral posterolateral thalamus (VPL) to the somatosensory cortex (S1) to form a conscious somatosensory percept ^9–11^.

Spatially arranged maps representing various sensory features have been discovered in the central nervous system, and they allow efficient processing of sensory information by grouping together relevant sensory, cognitive or spatial features. This probably allows the brain to more quickly and accurately process and respond to sensory information, potentially facilitating the organisation and integration of information from multiple senses and thus enabling more complex and sophisticated processing ^12^. One example hereof is the tonotopic layout of sound information in the auditory system. Here, sounds of similar frequencies are represented in spatially neighbouring regions in the brain ^13,14^. On the other hand, in the somatosensory system – which consists of a diverse group of LTMRs responding to a wide range of stimulation frequencies – it has been shown that adjacent areas of the body are represented by adjacent neurons. This has been termed somatotopy ^15^. However, it remains unclear if the somatosensory system uses a specific coding strategy to process frequency information, comparable to the tonotopy found in the auditory system.

To address this question we sought to characterise how both stimulus location and frequency are represented in individual neurons at different levels along the dorsal column-medial lemniscal pathway of the mouse hindlimb. Using two-photon calcium imaging, we revealed that overlapping somatotopic and tonotopic maps coexist in DCN, most likely reflecting heterogenous distribution of the receptor end-organ complexes in the periphery. These maps are partially preserved in VPL and S1. We thus propose that the central representations of vibrotactile stimuli combine somatotopic and tonotopic maps and form an integrated framework at earliest stages of information processing.

## Results

### Vibration frequency selectivity in somatosensory brainstem

To understand how information of fine discriminative touch is encoded in the central nervous system, we aimed to determine the functional organisation (somatotopy and tonotopy) of the brainstem DCN, where the first synaptic relay occurs in the somatosensory pathway (**Fig. 1a**). Pure sinusoidal vibrations of a wide range of frequencies (0.1 Hz to 3000 Hz, **Extended Data Fig. 1**) were delivered to the hindlimb of anaesthetised mice at various locations, from toe to thigh (see **Methods, Fig. 1a**). To characterise how vibrations are transmitted through the different parts of the hindlimb, we conducted non-invasive measurements with a laser Doppler vibrometer (**Extended Data Fig. 2**). These experiments demonstrated that the vibrotactile stimulation not only affects the skin locally (**Extended Data Fig. 2a**), but that it can be reliably conducted along the bones (**Extended Data Fig. 2b**). To measure the activity of neurons in the ipsilateral gracile nucleus, we expressed the fluorescent Ca^2+^ indicator GCaMP6s ^16^ in the medulla using viral vectors and imaged the responses under anaesthesia with two-photon microscopy through a glass window (**Fig. 1b**). We found that stimulation at the different locations or with different vibration frequencies, reliably evoked calcium responses, measured as transient increases in fluorescence of a subset of imaged neurons (**Fig. 1c, d, Extended Data Fig. 3**). This was consistent with the electrophysiological mapping of receptive fields in the previous literature ^17,18^. We then asked whether location or frequency are organised in a topographical manner (**Fig. 1e, f**). Surprisingly, we observed that location and frequency are both linearly organised along somatotopic and tonotopic axes, respectively (**Fig. 1g, h**). The experiments had been repeated in 6 mice and the results were reproducible (**Extended Data Fig. 3g, h**). Furthermore, these two map structures are consistent through the depth of the dorsal column nucleus (**Fig. 1i**), which is reminiscent of columnar architectures found in cortical maps ^15,19^. Lastly, we found that the gradients of the somatotopic and tonotopic maps are parallel (**Fig. 1j**).

**Figure 1.**
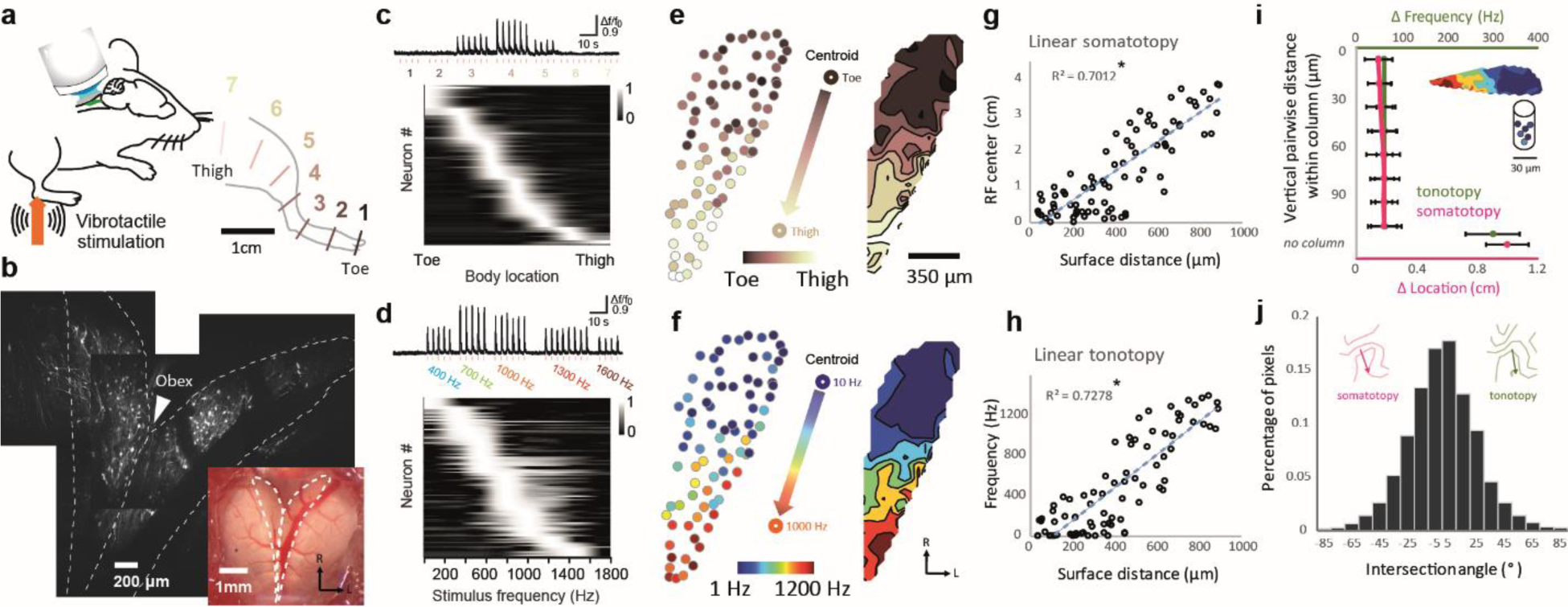
Maps for somatotopy and tonotopy in somatosensory brainstem, and their relationships. **a,** Experimental setup schematic. Hindlimb of the individual mouse, from toe to thigh, is divided into seven locations for delivery of vibrotactile stimuli for location preference mapping. **b,** Dorsal view of the imaging window above dorsal column nuclei (DCN) showing the expression patterns of GCaMP6s. White dashed lines indicate the boundaries of the left and right gracile nuclei. The white arrowhead points the obex and the midline **c.** Top, relative changes in fluorescence (Δf/f0) of an example neuron imaged with two-photon microscopy in response to stimulation (100 Hz vibration at an amplitude of 400 µm) in seven different locations. Bottom, normalised tuning curves for all neurons in one animal sorted by preferred location (n/N = 98/1). **d,** Top, Δf/f0 of the same example neuron in response to vibration at its preferred location in different frequencies. Bottom, normalised tuning curves for all neurons in one animal sorted by decreasing preferred frequency (n/N = 98/1). **e-f,** Left, each neuron’s preference for location (**e**) and frequency (**f**), colour-coded at their physical position in DCN. Middle, somatotopic (**e**) and tonotopic (**f**) axes defined by the centroid of the responsive populations (See **Methods**), Right, preference maps of somatotopy (**e**) and tonotopy (**f**) interpolated from the imaged neural population. **g-h,** The relationship between surface distance of the nucleus in the DCN and location (**g**) or frequency (**h**) along the somatotopic axis from toe to thigh (n/N = 98/1, linear regression, all pairs P > 0.05). **i,** The relationship of vertical distance between neurons in a pair within a column, and their difference in location (pink) and frequency (green) is calculated (n/N = 892/7, Kruskal–Wallis test, P < 0.0001). Error bars represent S.E.M. **j,** Histogram showing the distribution of the intersection angle between gradients of somatotopic map and tonotopic map. A circular Rayleigh test showed that the distribution was significantly different from uniform (n/N = 892/7, Circular Rayleigh test, P < 0.0001).

### Maps for somatotopy and tonotopy in somatosensory pathways

Where else along the somatosensory pathway of mice could this kind of topographical organisation be observed? To address this question we performed two-photon calcium imaging in all hubs of the dorsal column-medial lemniscal pathway (**Fig. 2 a-d**). Here, we analysed the pairwise distance between neurons and their difference in preferred location or frequency, without presumption of a linear topographical organisation. Based on the relationship of horizontal distance between paired neurons and their difference in the selected features, we did not observe fine-scale topological organisation for either location or frequency in the lumbar dorsal root ganglia (DRG), but rather a salt and pepper-like organisation (**Fig. 2e, Extended Data Fig. 4**). In contrast, we found that in the gracile nucleus of the DCN, the relationship between the change in physical distance and the average change in preferred feature is extended over a few hundred micrometres (**Fig. 2f**), consistent with the linear organisation described in Figure 1. We, next, implanted a GRIN lens directly over the ventral posterolateral nucleus of thalamus. These recordings revealed that the fine-scale somatotopy and tonotopy was still present at the level of the thalamus (**Fig. 2g**), though less pronounced compared to the brainstem, as the significant difference between the original and the shuffled data could only be observed within 25 μm pairwise distance. Finally, the two fine-scale topological organisations were still partially preserved in the hindlimb somatosensory cortex, which could also be visualised from the slope of the linear fit (**Fig. 2h**).

**Figure 2.**
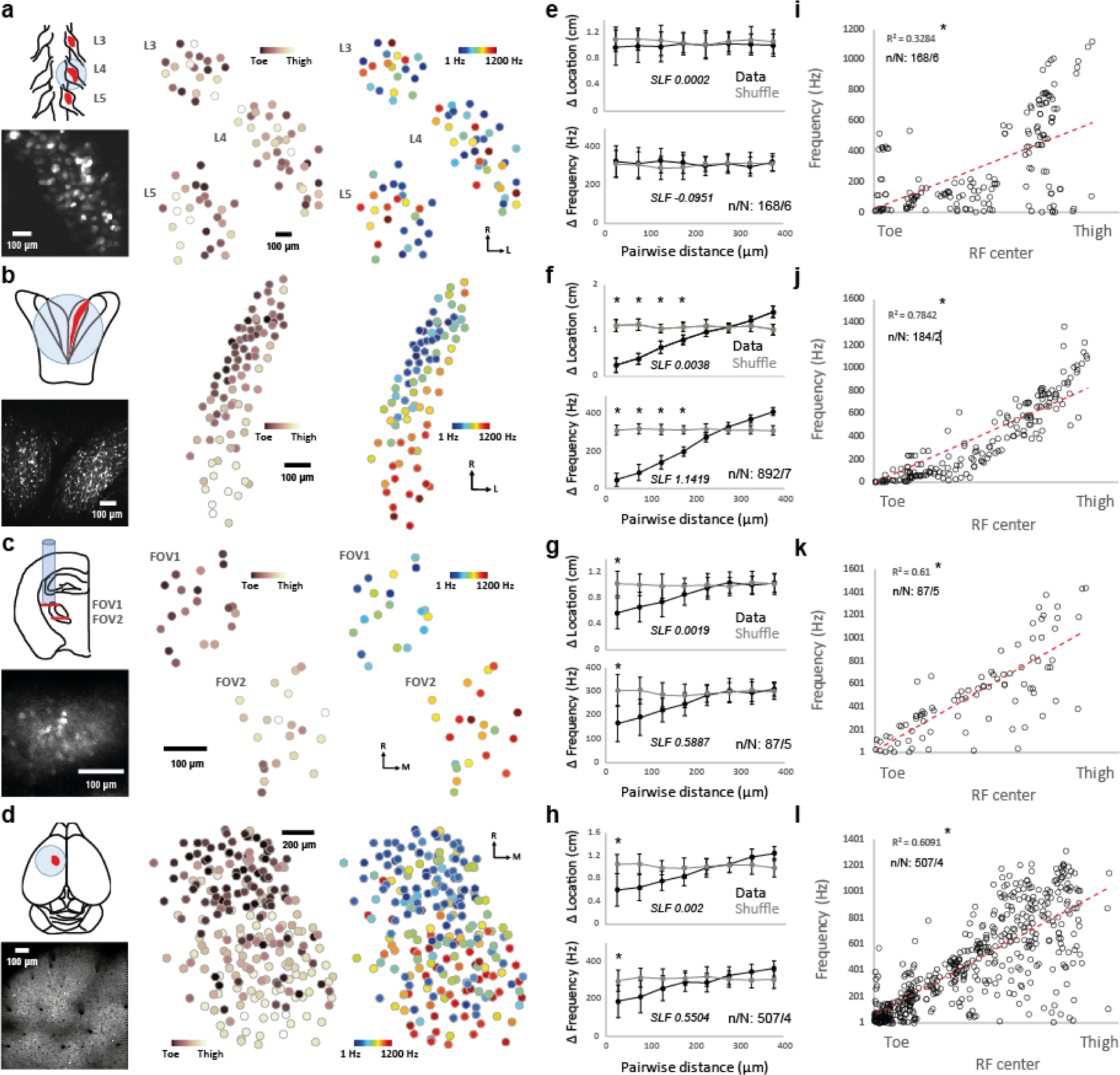
Maps for somatotopy and tonotopy in somatosensory pathways for discriminative touch. **a-d,** Left, schematic of the clusters of neuronal cell bodies (top) and example fields-of-view (bottom) along the somatosensory pathway, which ascends from the lumbar dorsal root ganglia (**a**), to the gracile nucleus of the DCN (**b**), then the ventral posterolateral nucleus of thalamus (**c**), and finally the hindlimb somatosensory cortex (**d**). Right, each neuron’s preference for location and frequency is colour-coded. **e-h,** The relationship of horizontal distance between neurons in a pair and their difference in location (top) and frequency (bottom) is calculated (Permutation test, *P < 0.01) for lumbar dorsal root ganglia (**e**), gracile nucleus (**f**), ventral posterolateral nucleus of thalamus (**g**), and hindlimb somatosensory cortex (**h**). The slope of the linear fit (SLF, italic) of the first 5 data points from the original data (black line) is shown. Error bars represent S.E.M. **i-l,** The correlation of location and frequency of individual neurons (linear regression, *P < 0.01) for lumbar dorsal root ganglia (**i**), gracile nucleus (**j**), ventral posterolateral nucleus of thalamus (**k**), and hindlimb somatosensory cortex (**l**), showing the joint representation of location and frequency changed dramatically.

Furthermore, we investigated the transformation of the joint representation of location and frequency along the somatosensory pathway. At the peripheral level, the relation between location and frequency presents as a weak, but significant, linear correlation (**Fig. 2i**). This result is consistent with the distribution of mechanosensitive end organs in mouse hindlimb (**Extended Data Fig. 5**), implying that the tonotopic map might be partially inherited from the distribution of mechanosensitive end organs ^20^. Although there is a correlation between location and frequency in all four neuronal structures, significant changes of the combined representation could be observed between periphery to brainstem and between brainstem to thalamus, from a low correlation (DRG) to a high correlation (DCN) and back to a moderate correlation (VPL, S1), respectively (**Fig. 2i-l**, two-dimensional Kolmogorov-Smirnov test, DRG-DCN, p = 0.0431; DCN-VPL, p = 0.0055; VPL-S1, p = 0.3594). Overall, we speculate that the emergence of the fine-scale tonotopy occurs at the level of the DCN, and thus between the peripheral and central nervous systems, where the synaptic integration from various mechanoreceptors for gentle touch happens for the first time in the dorsal column-medial lemniscal pathway.

### Peripheral origin of somatotopic and tonotopic maps

In order to understand how the convergence of the different peripheral mechanosensitive end organs could contribute to the emergence of the tonotopic map in the brainstem, we aimed to first characterise the threshold profile of each major end organ type in the mouse hindlimb. Here, we performed *in vivo* nerve fibre recordings ^17,21^ (see **Methods**) of the sciatic nerve in anaesthetised mice. Ramping stimulations with a wide range of frequencies (0.1 Hz to 3000 Hz) were applied to define the response threshold for each mechanosensitive afferent (**Fig. 3a, Extended Data Fig. 6**). The threshold curve of each individual fibre was mapped and then pooled into one of the three threshold curves (**Fig. 3b**). In accordance with the literature ^3^, each of these three groups can be matched with the physiological properties of the LTMR likely innervating Merkel cells (Aβ SA-LTMR), Meissner corpuscles (Aβ RA1-LTMR) and Pacinian corpuscles (Aβ RA2-LTMR); for simplification, we will use the acronyms, MERK, MEIS and PACI to indicate their response properties and link them to their typical receptor end organs.

**Figure 3.**
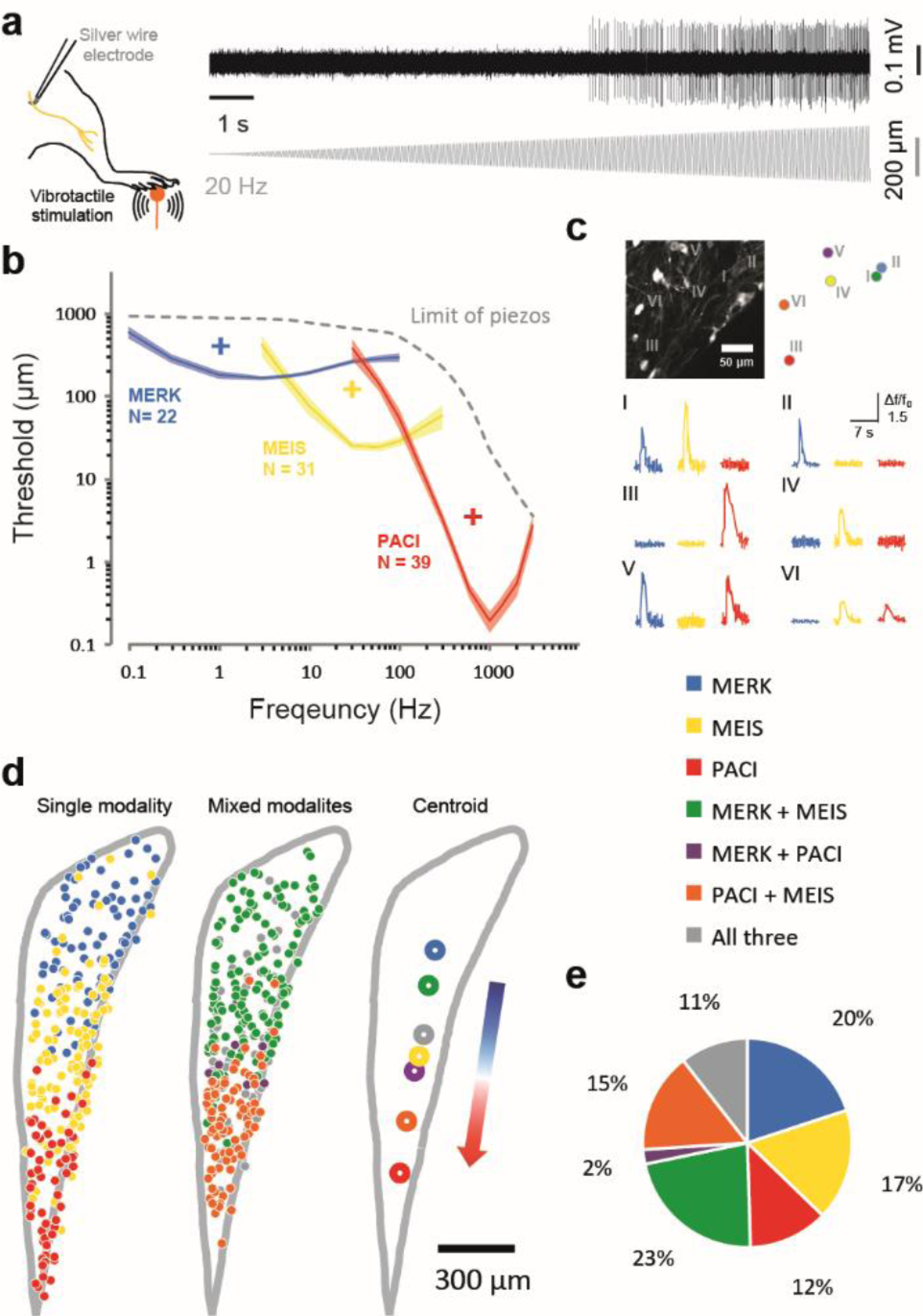
Convergence of peripheral mechanosensitive end organs in the brainstem. **a,** Left, schematic of *in vivo* nerve fibre recording of sciatic nerve. Right, representative trace of a mechanosensitive afferent spiking rate in response to 20 Hz vibration stimulus. **b,** The averaged threshold curves from three major groups of mechanosensitive end organs (n/N = 92/10). Shaded area indicates S.E.M. +, the stimulation parameter specific to preferentially activate only one defined group of mechanosensitive end organs: Merkel cell-like (MERK), 1 Hz, 300 μm; Meissner-like (MEIS), 30 Hz, 100 μm; Pacinian-like (PACI), 600 Hz, 3μm. **c,** Top, an example field-of-view (left) and colour-coded response identities (right) of neurons in gracile nuclei. Bottom, example traces of the six neurons responding to stimulation derived from **b**. **d,** Color-coded response identities of neurons in gracile nuclei across animals (n/N = 477/4). Left, the centroid of each responding group. Quantification of the spatial distribution of neurons was depicted relative to the centroid of Merkel cells-specific response and to Pacinian corpuscles-specific response (see methods). **e,** The percentage of seven classified groups shows around half of the population responds to more than one mechanosensitive end organs-specific stimulation. Number of experiments and statistics in Extended Data Table 1-2.

The clear separation of frequency tuning observed in the mouse can be used as a basis to determine the stimulation parameter specific to activate only a unique group of mechanosensitive end organs. We thus applied three end organ-specific stimuli (plus signs in **Fig. 3b**) to the hindlimb while measuring the stimulus-related activity in DCN (**Fig. 3c**). We discovered nearly half of the population can be activated by more than one type of stimulation, indicating that the suprathreshold level of convergence from more than one end organ type at the single DCN neuron is prevalent (**Fig. 3e**). Subsequent calcium imaging of the dendrites of individual DCN neurons further revealed that the subthreshold level convergence can occur at the level of individual spines (**Extended Data Fig. 7**). Moreover, the spatial distribution of the neurons receiving inputs from different types of end organs (**Fig. 3d**) is also consistent with the linear pattern of tonotopy (**Fig. 1f**) with the difference of vector angles smaller than 10°, implying a potential underlying axonal organisation serving as the foundation of synaptic integration.

### Synaptic mechanism underlying the emergence of tonotopic map for substrate vibration

To explicitly test the hypothesis that the axonal afferents of the LTMRs innervating differently tuned end organs would preferentially target specific areas of the DCN according the tonotopic organisation ^7,22^, we expressed calcium indicators in the lumbar spinal dorsal horn and DRG (see **Methods**) and imaged the activity in their axonal terminals within the gracile nucleus (**Fig. 4a, Extended Data Fig. 8**). After combining data from multiple animals (N = 7), the clustering analysis revealed a systematic distance-preference relationship of the boutons in DCN (**Fig. 4b**), similar to the cellular level of somatotopy and tonotopy (**Fig. 2f**). We next wanted to understand how this organisation was transferred to the postsynaptic networks of gracile nucleus neurons; we thus sparsely expressed the glutamate sensors iGluSnFR3 ^23^ in the DCN (**Fig. 4c, Extended Data Fig. 7**). A similar functional organisation was observed among the spines of excitatory synapses of the DCN neurons (**Fig. 4d**), reflecting the somatotopic and tonotopic features described in the presynaptic networks. On top of that, local clusters of inputs encoding similar location and frequency could also be found in the short segment of the dendrites, which might contribute to the non-linear dendritic integration (**Extended Data Fig. 7**). Lastly, we wanted to test the hypothesis of whether simply the random sampling of presynaptic networks innervated from lumbar spinal level can explain the emergence frequency selectivity of individual DCN neurons. For this purpose, we used the averaged tuning curves from either the actual synapses of the imaged neuron or the random sampled synaptic inputs from the boutons within the dendritic field as a predictor of its somatic tuning. We found that in the case of random sampling, the predictability of somatic tuning cannot reach that of the actual synaptic inputs (**Fig. 4e-f**). This simulation supports the idea that the functional specificity of connections, in other words selective dendritic sampling, refines the frequency tuning of individual neurons within the tonotopic map.

**Figure 4.**
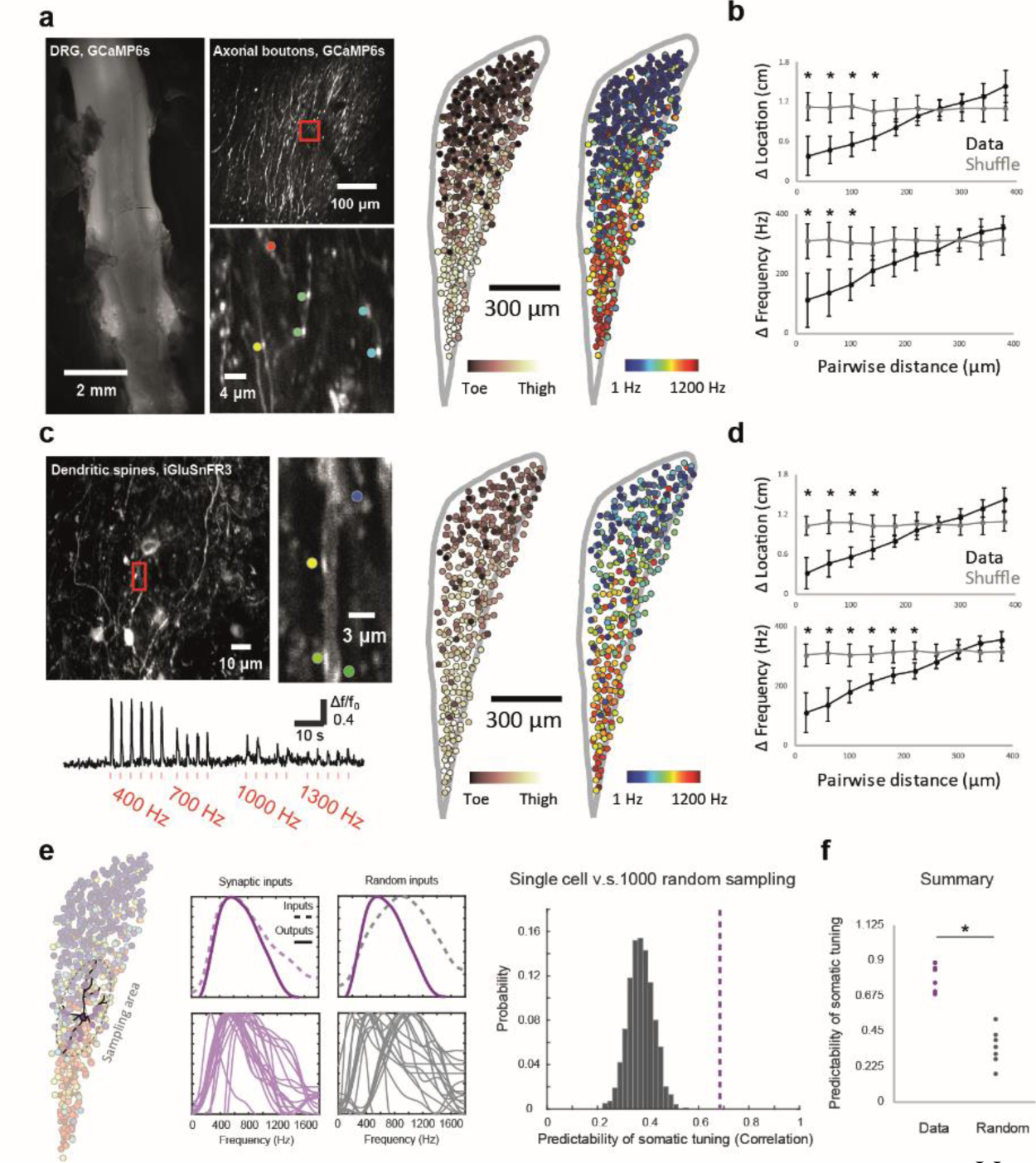
Functional specificity of connections within dendritic fields of brainstem neurons. **a,** Left, dorsal view of lumbar segment of spinal cord (L3-L6) and dorsal root ganglia expressed by GCaMP6s, overall field-of-view in gracile nucleus (upper right), and the individual imaged field-of-view of axonal boutons (lower right). Right, each bouton’s preference for location and frequency colour-coded. **b,** The relationship of distance between boutons in a pair and their difference in location (top) and frequency (bottom) is calculated (n/N = 913/7, Permutation test, *P < 0.01). Error bars represent S.E.M. **c,** Left, overall field-of-view of sparse neurons expressed iGluSnFR3 in gracile nucleus, and the individual imaged dendritic segments. Right, each bouton’s preference for location and frequency colour-coded. **d,** The relationship of distance between synapses in a pair and their difference in location (top) and frequency (bottom) is calculated (n/N = 410/6, Permutation test, *P < 0.01). Error bars represent S.E.M. **e,** Left, an example neuron with reconstructed dendritic field and synaptic inputs, overlaying on the bouton frequency preference scatter plot from **c**. Middle, the tuning curves of 20 actual synaptic inputs (lower left) and predictability of the somatic tuning curve based on their average (upper left); the tuning curves of 20 random sampled synaptic inputs from the boutons within the dendritic field (lower right), and their predictability of the somatic tuning curve (upper right). Right, the comparison of predictability between random sampling and actual synaptic inputs. **f,** The summary results of functional specificity of connections from 7 neurons (n/N = 7/5, Wilcoxon rank-sum test, *P < 0.01).

## Discussion

Traditionally, the somatosensory brainstem is viewed as a simple relay for sensory information from the periphery to the higher brain regions. In this study, we show that the dorsal column nuclei are a crucial integration site where at least two parallel and gradual topographical maps coexist along the rostral-caudal axis, one representing the stimulus location along the limbs, the other representing the different vibration frequencies. In addition, we observed that these maps not only reflect the distribution of mechanosensitive end organs in the periphery, but also the neural circuits in the brainstem. Whereas the overall somatotopic organisation of brainstem nuclei have been described in numerous mammalian species ^7^, the recent discovery of a cellular-scale brainstem map for visceral sensations suggests this may be a common strategy for the central nervous system to encode the sensory information from the periphery ^24^ at the level of the brainstem. In our study, we now show that additional computational layers exist in parallel with the classical somatotopy. This might reflect a way to optimise for uniform coverage of multiple features ^25–27^. Though it has been described that the gracile in rats, cats and primates is subdivided into substructures ^7^, we didn’t observe any clear organisation in the mouse in neither in vivo imaging nor histology.

The well documented tonotopy in the auditory system, is proposed to play an important role in allowing the brain to process and interpret auditory information efficiently and effectively, which is crucial for our ability to communicate, understand speech, and appreciate music ^13^. What could be the functional role of a tonotopic organisation of somatosensory information we reveal here? We can speculate along several lines: on the one hand, the exquisite sensitivity (∼0.01 μm) for vibrotactile stimuli and the robust high frequency responses (∼3 kHz) might represent an extension of the auditory system of the mouse ^28^. The range of lowest perceptible vibrotactile responses (0.1-3 kHz) ideally complements their range of auditory frequencies (∼3-60 kHz, ^29^). A tonotopic representation of vibrotactile frequencies would thus allow for a similar type of processing compared to the auditory system and could be part of the neural basis for the efficient discrimination of vibrotactile pitch ^28^. On the other hand, one can speculate that a gradual tonotopic organisation might be ideally suited to process relative frequency information and participate in the normalisation process required during active sensation ^30,31^. For example, a surface texture is perceived the same way, independent of the speed with which it is explored. More detailed analysis will be required to address these questions in future experiments. Finally, we found that the biomechanics of the hindlimbs show a rather robust conductance of all frequencies along the bones and joints (**Extended Data Fig. 2**), which suggests that the Pacinian corpuscle located near the bone can probably sense the vibration from any stimulus that successfully reaches the fibula. In contrast, vibration applied to the distal paw regions tended to attenuate rapidly, suggesting a more discrete system.

In addition to the overlapping functional maps, we also demonstrate that the information from different mechanoreceptor end organs is heavily integrated at the level of the DCN (**Fig. 3**) and thus forms a continuous tonotopic representation at the earliest possible stage (**Extended Data Fig. 9**). This direct dendritic mapping and detailed dendritic computation at the brainstem level ^18,32^ is reminiscent of the dendritic computation that underlies the emergence of orientation selectivity in the visual cortex ^33,34^ and might thus represent a general principle for the emergence of such maps across sensory modalities. Another remaining challenge will be to dissect the contribution of individual mechanosensory units, such as by optogenetically targeting the LTMRs that innervate Meissner and Pacinian corpuscles, and Merkel cells. Several genetic targeting or intersectional methods with viral injections have been developed to selectively activate sensory afferents in the periphery ^11,35^, but end-organ specificity remains challenging. Although the DRG neurons are the main inputs to the DCN, it would also be interesting to explore other long-range excitatory inputs, local or feedback inhibition which might jointly shape the output of DCN and thus the outcome of the perception and behaviour ^30^. Consistent with the results of recent studies ^9,36^, we confirmed by pharmacological inactivation of the spinal dorsal horn (**Extended Data Fig. 10a-b**), that the indirect dorsal column pathway does not contribute significantly to the encoding of high frequency vibration in the DCN. The block of excitatory synaptic transmission within the spinal cord only affected the response amplitudes of low frequency selective DCN neurons (**Extended Data Fig. 10c**).

Finally, we hope our data will also help to constrain network models on the mechanisms of neuronal tuning in all the output regions of DCN, including thalamus, spinal cord, cerebellum, tectum, and other brainstem nuclei ^7^, which are usually intimately linked to the emergence of new feature selectivities and their corresponding maps ^37^, and are ultimately the basis for the conscious perceptions of vibrotactile cues.

---

**Extended Data Figure 1.**
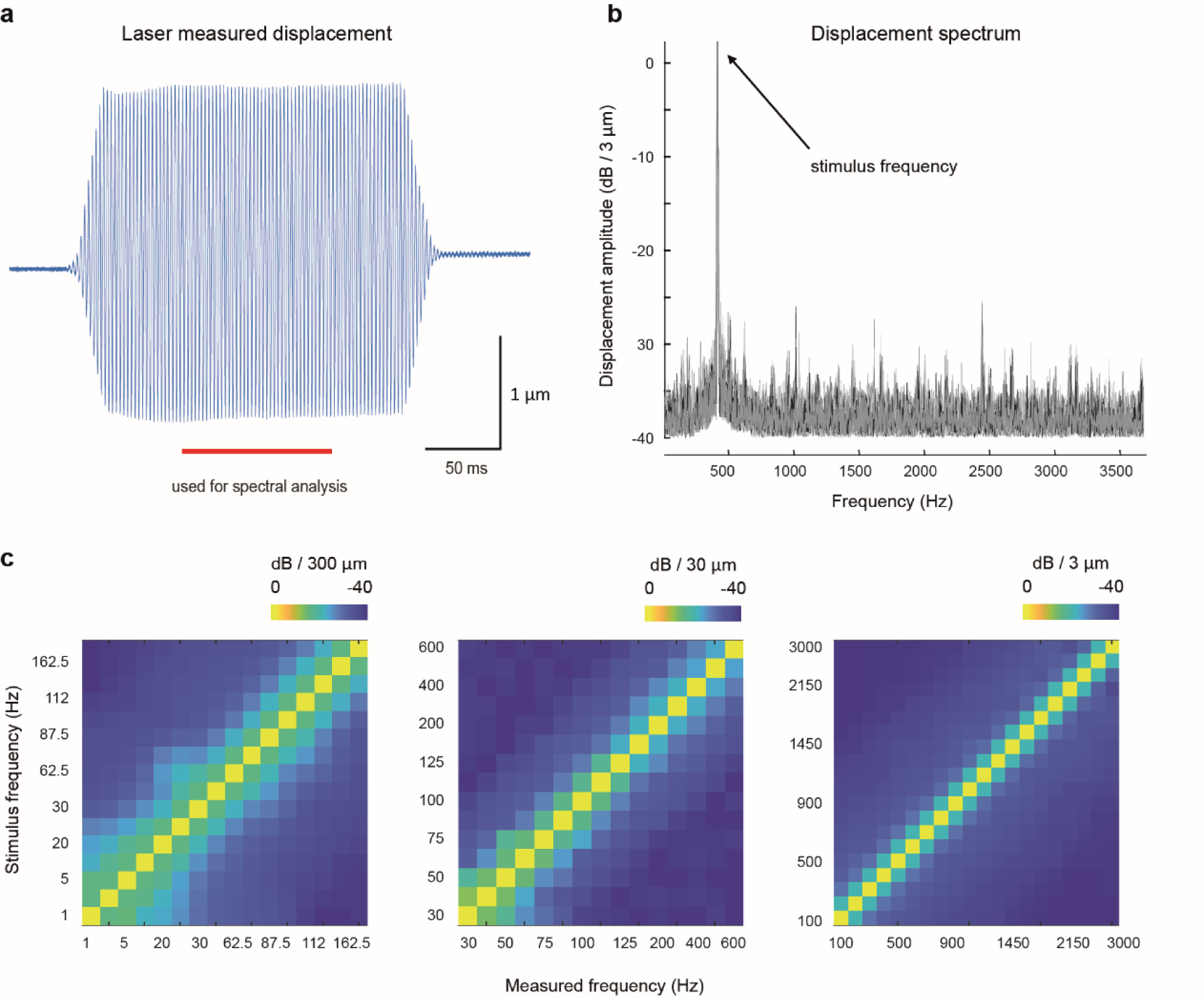
Spectral analysis of vibrations stimulus measured with a laser doppler vibrometer. **a,** Example of a displacement measured by the laser doppler vibrometer of a single 400-Hz vibration. The middle-half interval of spectral analysis was highlighted in red. **b,** Amplitude spectrum of the displacement produced by 10 trials of 400-Hz stimuli indicates that the physical vibration consisted of a single frequency component (nearly a pure sinusoid). Components apparent at the distant frequencies were highly attenuated relative to the stimulation frequency, similar to the background noise. **c,** Amplitude spectra were obtained for stimuli at all tested frequencies and three representative amplitudes from two different piezo actuators.

**Extended Data Figure 2.**
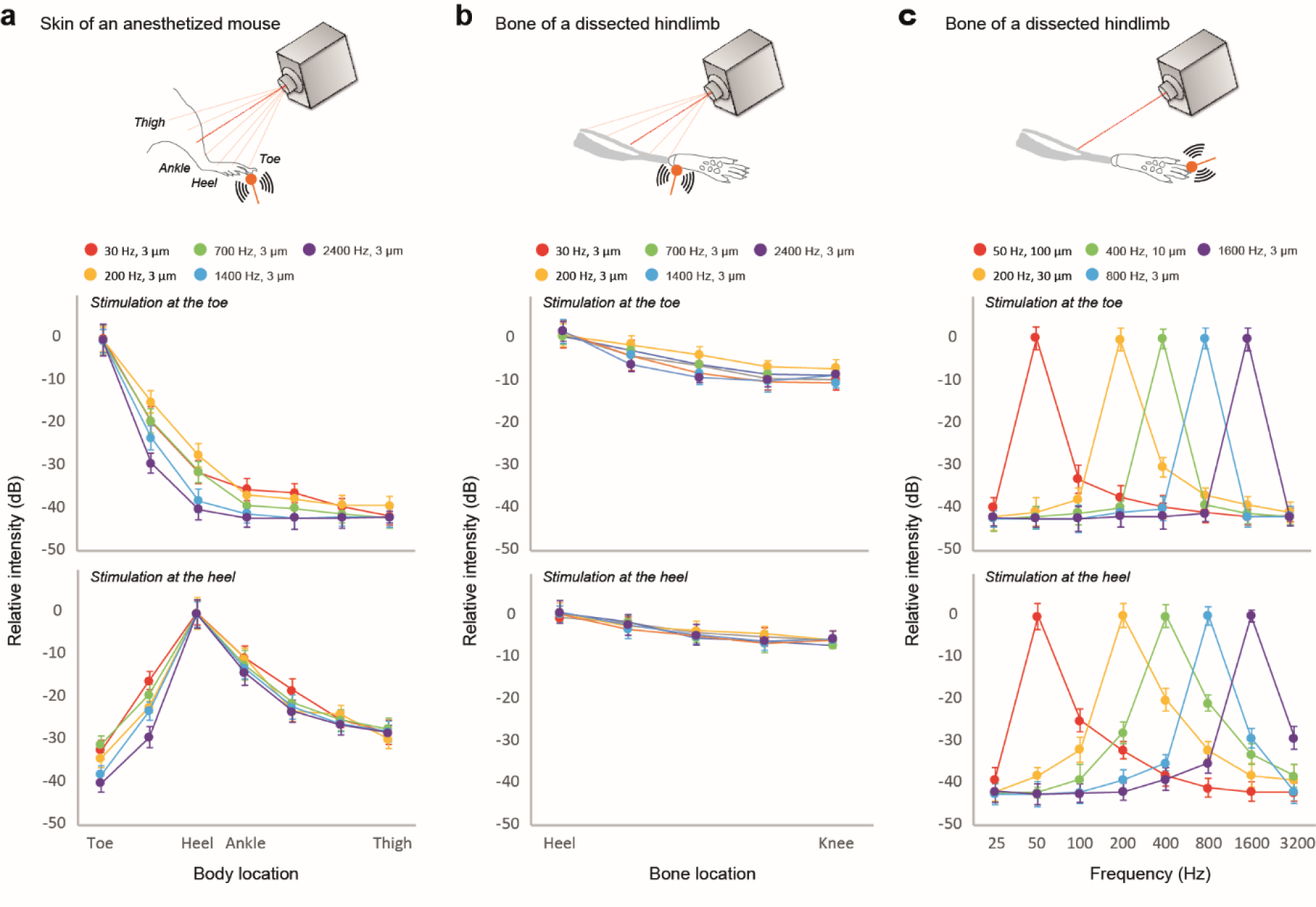
Transmission of vibration within the hindlimb tissue. **a,** Amplitude spectra measurement of transmitted vibration along the 7 locations of the hindlimb of an anaesthetised mouse with a laser doppler vibrometer (N = 2). Top, stimulation was placed at the toe. Bottom, stimulation was placed at the heel. **b,** Amplitude spectra measurement of transmitted vibration along the 5 along the fibula freshly dissected from an euthanized mouse. Top, stimulation was placed at the toe. Bottom, stimulation was placed at the heel. **c,** Signal intensity of different frequency components were obtained *ex vivo* at the base of fibula in the dissected hindlimb, where the Pacinian corpuscle mainly located. Top, stimulation was placed at the toe. Bottom, stimulation was placed at the heel. Error bars represent S.E.M.

**Extended Data Figure 3.**
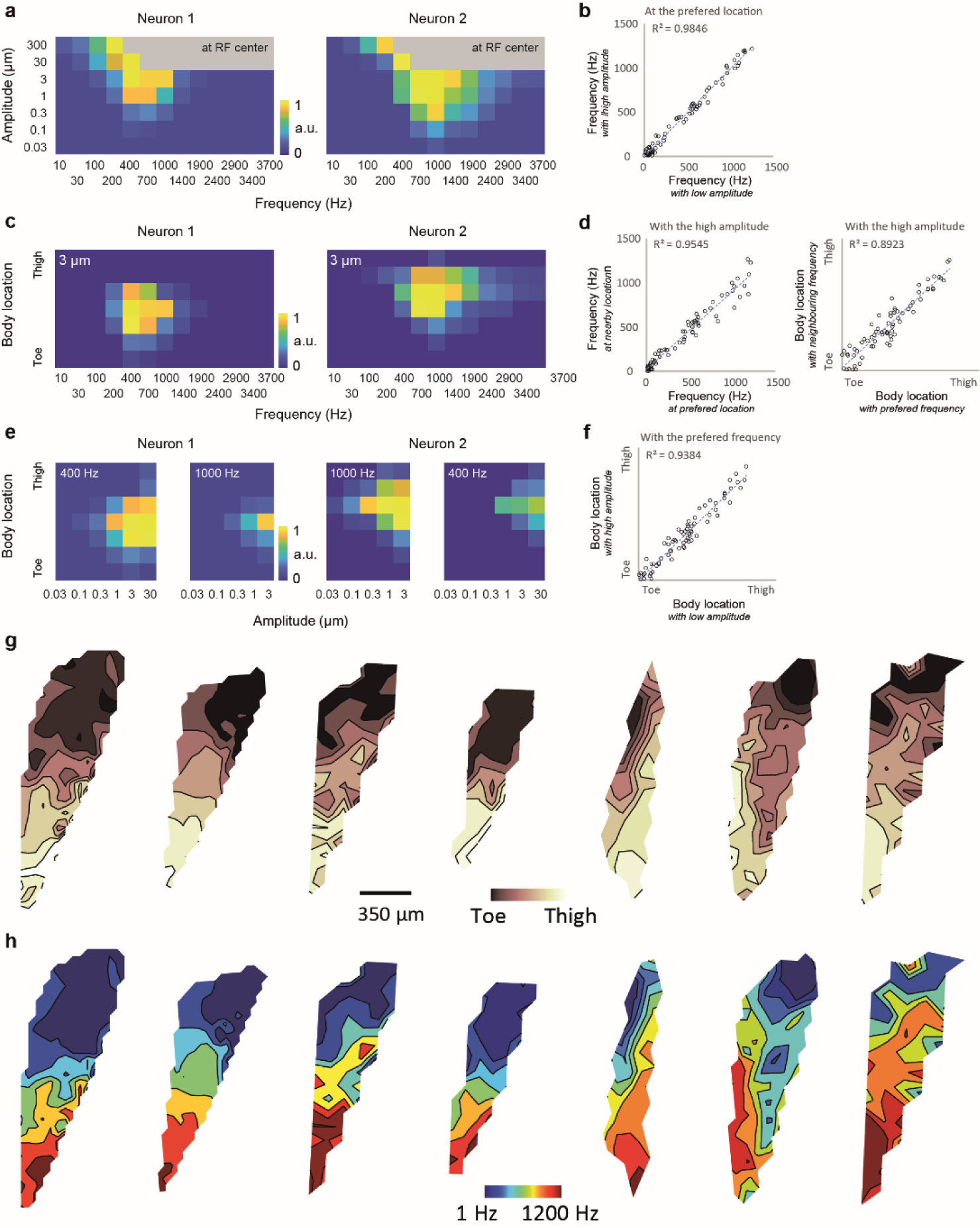
Complete DCN preference maps, invariance of the receptive field properties and the interaction between frequency, location and amplitude. **a,** Joint selectivity of the vibration frequency and the amplitude from two DCN neurons, with stimulation applied to the receptive field centre. Note, that the high frequency stimulation of pure sinusoidal vibration cannot be generated at the high amplitude due to the technical limits. **b,** The correlation of the frequency preference of individual neurons measured with highest and lowest vibration amplitudes, which yielded a completed tuning curve. **c,** Joint selectivity of the frequency and the location from the two neurons, with the highest stimulation amplitude that yielded a completed tuning curve. **d,** Left, the correlation of the frequency preference of individual neurons measured at the different body location, one at the receptive field and one at the nearby location with the second best responses. Right, the correlation of the location preference of individual neurons measured with different frequencies, one with the preferred frequency and one with the second-best preferred frequency. **e,** Joint selectivity of the location and the amplitude from two DCN neurons, with two frequencies that highly activated the neurons. **f,** The correlation of the location preference of individual neurons measured with highest and lowest vibration amplitudes, which yielded a completed tuning curve. Number of experiments and statistics in Extended Data Table 1-2. **g-h,** The DCN preference maps of somatotopy (**g**) and tonotopy (**h**) from all seven animals. The first one is the same as the example in Figure 1.

**Extended Data Figure 4.**
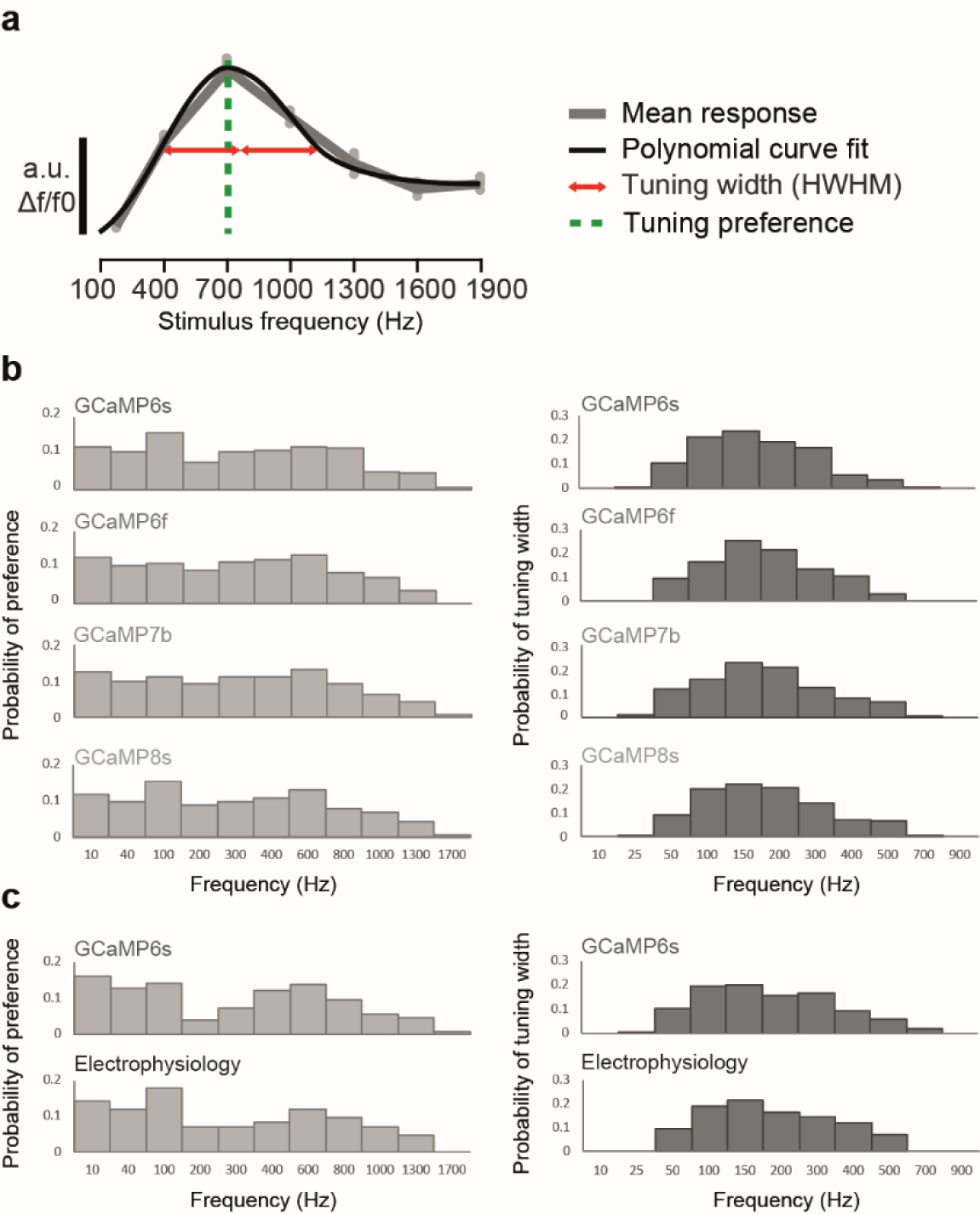
Comparing different generations of GCaMP imaging and electrophysiology. **a,** The example tuning curve of a DCN neuron across different vibration frequencies at the vibration amplitudes of 3 µm: grey dots, individual stimulus responses; grey trace, mean values; black trace, polymodal function fits; vertical green dashed line, preferred frequencies; red horizontal lines, the half-width at the half-maximum of the curve. **b,** Left, the distribution of the tuning preference from the DCN imaging experiments performed with different generations of calcium indicator, GCaMP. Right, the distribution of the tuning width measured in the different imaging experiments. **c,** Left, the distribution of the tuning preference measured from the calcium imaging at the DRG (top) or electrophysiology recording at the nerve fibres (bottom). Right, the distribution of the tuning width measured from the calcium imaging (top) or electrophysiology experiments (bottom). No significant difference was found in any comparison within DCN or DRG. Number of experiments and statistics in Extended Data Table 1-2.

**Extended Data Figure 5.**
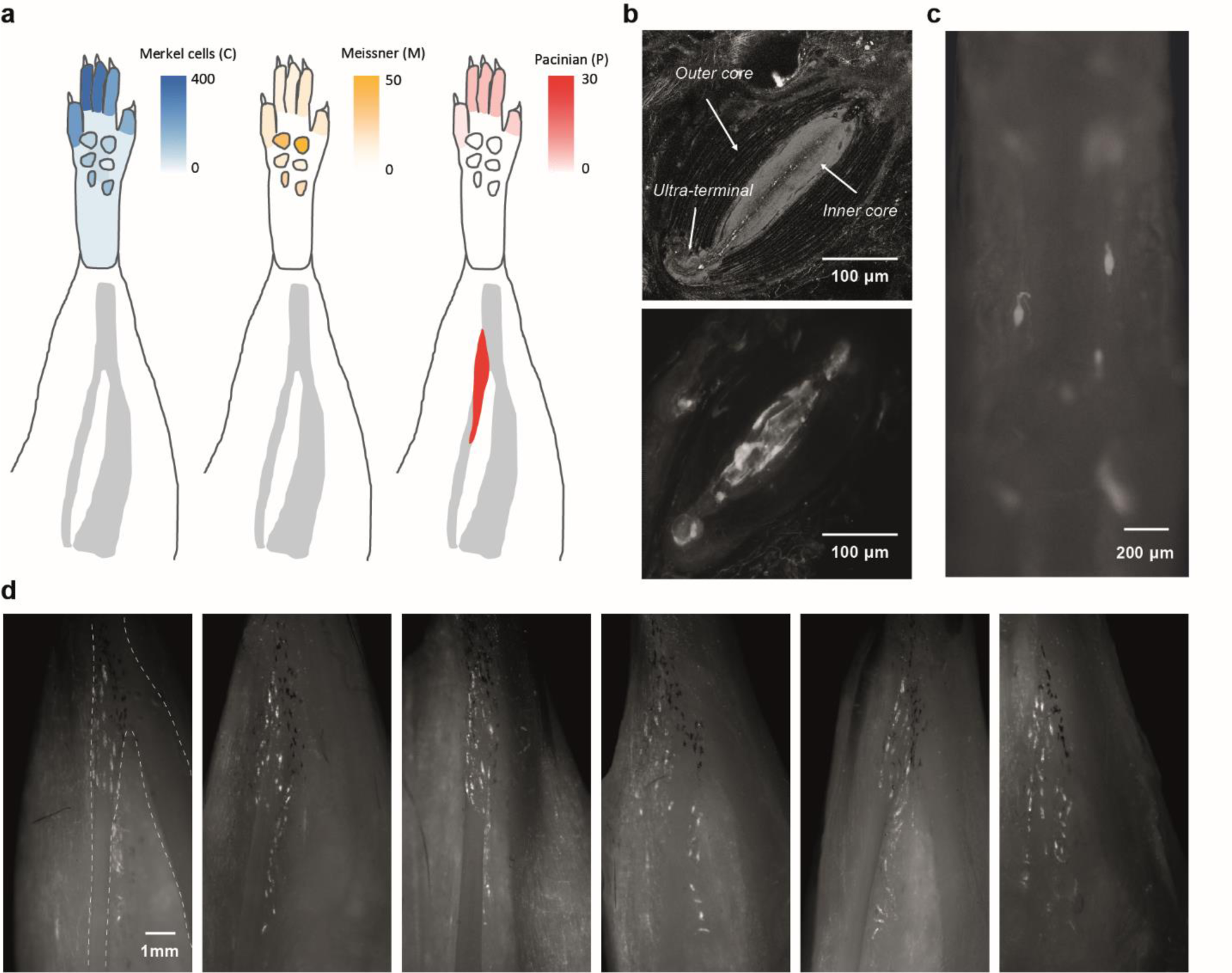
Anatomical distribution of mechanosensitive end organs in hindlimb. **a,** Summary of the distribution of the three main mechanosensitive end organs in the mouse hindlimb (n/N = 243/5). Data of Merkel cell and Meissner corpuscle distribution in the paw is interpreted from the literature ^20^. There was no quantitative data for the distribution of the Merkel cells and Meissner corpuscles beyond the hindpaw. **b,** Top, a Pacinian corpuscle from a wild-type mouse under the confocal imaging after dissection *ex vivo*. The denser autofluorescence signals from the inner core cells and the onion-like layers of outer core cells could be visualised with high laser power and gain of the photon detector. Bottom, the inner core cells of a single Pacinian corpuscle were visualised using a double-transgenic strategy (ETV1-Cre x tdTomato) under *in situ* two-photon imaging at the hindlimb. **c,** Example plantar field of view of the toe with removed skin, where two tdTomato-positive Pacinian corpuscles are found close to the phalange. **d,** Six example hindlimbs with skin and muscle removed to expose the distal tibiofibular joint, and they consistently showed many tdTomato-positive Pacinian corpuscles (each bright spot is ETV1+ cells in the inner core area of one Pacinian corpuscle) wrapping around the fibula bone.

**Extended Data Figure 6.**
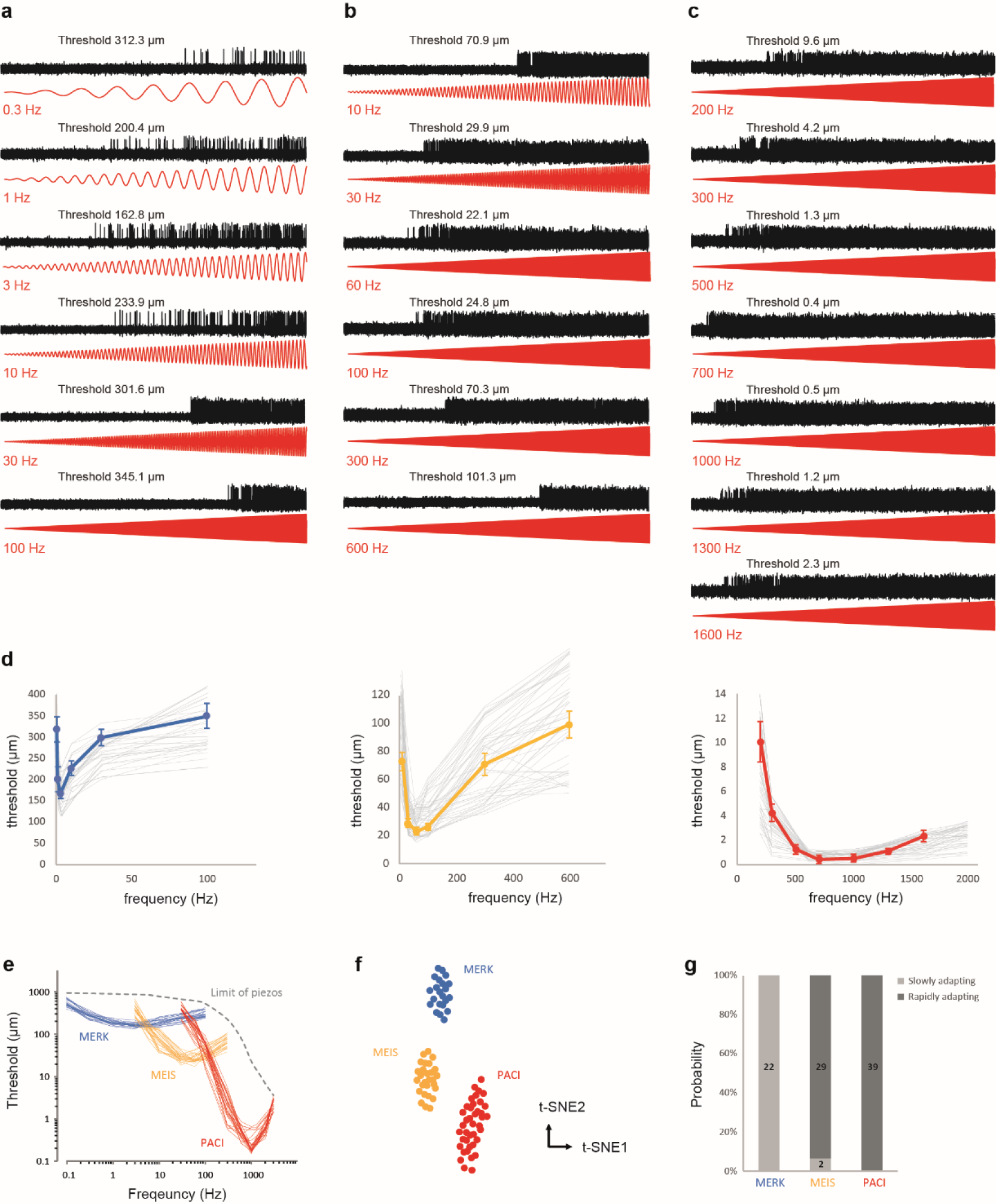
Details of the threshold mapping of mechanosensitive afferents. **a-c,** Mechanosensitive afferents were stimulated by applying a series of sinusoidal mechanical stimuli. These were presented at the receptive field centre of individual units,at linearly increasing amplitude and with increasing frequencies (0.3–1600 Hz). Examples of responses evoked by full range of the stimulation of mechanosensitive afferents potentially innervating Merkel cells (a) and Meissner (b) and Pacinian corpuscles (c), respectively. The threshold was determined by the first spike. **d,** Frequency dependent threshold curves (mechanical activation threshold versus stimulation frequency) of the three mechanosensitive afferents above. Data represent the mean ± S.E.M. The grey lines show the mean threshold curve of other nerve fibres. The exact frequencies used for mapping were different for individual nerve fibers in some experiments. **e,** The threshold curves of all three groups of nerve fibres with a logarithmic scale. **f,** Functional clusters shown in a t-SNE space. **g,** Adaptation characteristics were defined by if there were any action potentials in response to a static indentation. Number of the mechanosensitive afferents was indicated.

**Extended Data Figure 7.**
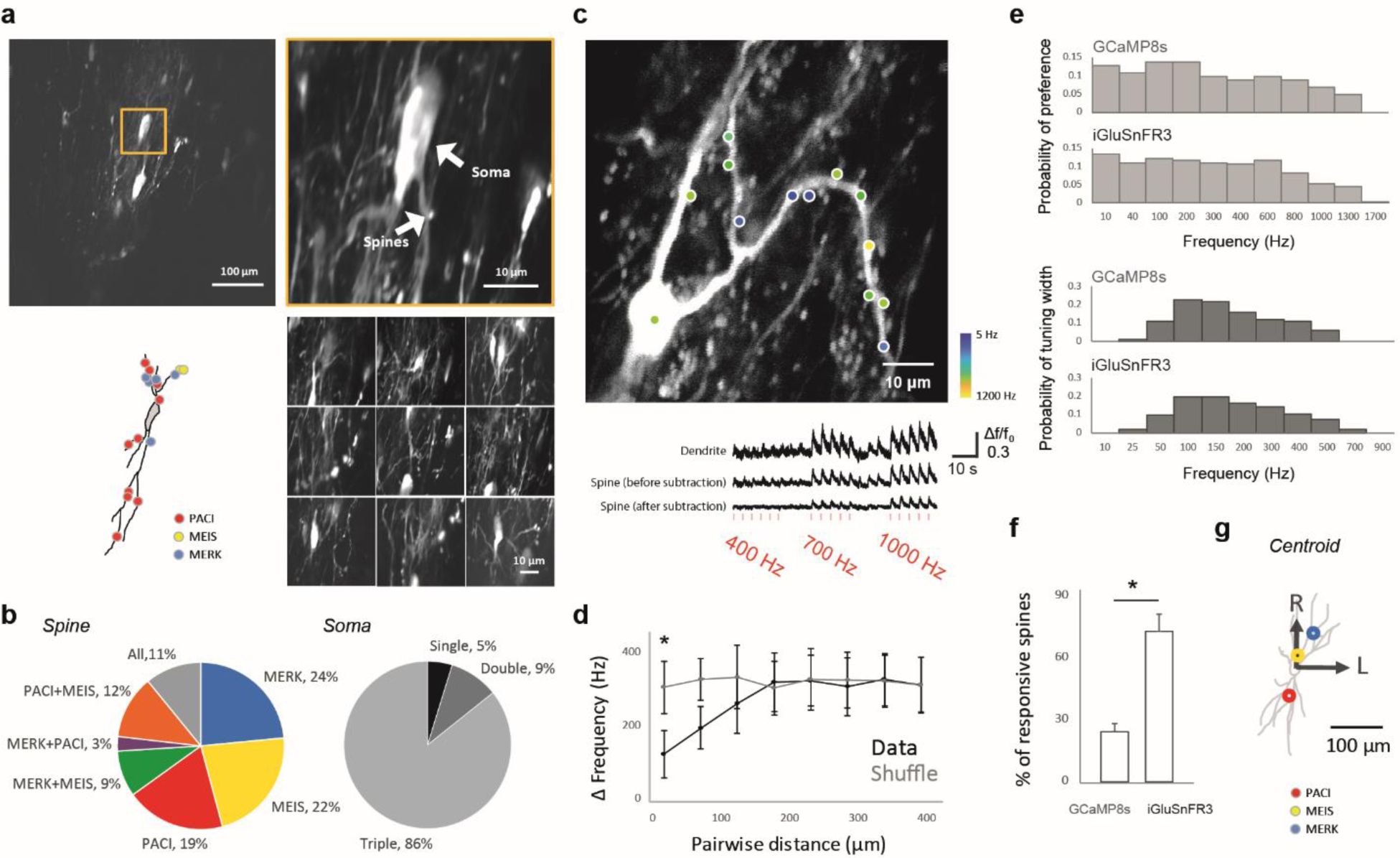
Details about dendritic spine calcium and glutamate imaging. **a,** Convergence between different end-organ types in the dendrites of single DCN neurons. *In vivo* two-photon calcium imaging of a DCN neuron with its dendritic field (top left). An example field of view with the soma of the targeted cell (top right). Nice example fields of view covering a part of the dendritic arbour (bottom right). Functional organisation of various submodalities in the dendritic field of the targeted cell (bottom left). **b,** Left, the percentage of seven classified groups of all dendritic spines in DCN defined by three mechanosensitive end organs-specific stimulation. Right, the percentage of DCN neurons receiving synaptic inputs responding to different numbers of stimulation (18 out of 21 neurons are with spines responding to all three stimulations, 2 neurons are with spines responding to at least two types of stimulations, and just 1 neuron is with spines belonging to response properties MEIS). **c,** Top, the distribution of different frequency preferences (tonotopy organisation) on a dendritic segment of the DCN neuron labelled with GCaMP8s. Bottom, postsynaptic activity from a single example spine was isolated by subtracting the component of the signal from the dendritic shaft. Red line: Duration of light stimulus. **d,** The relationship of distance between spines in a pair and frequency preference was calculated and it revealed that spines with a similar orientation preference were more likely to cluster. Error bars represent S.E.M. *P < 0.001, permutation test. **e,** Top, the distribution of the tuning preference of spines measured from the GCaMP8s calcium imaging or iGluSnFR3 glutamate imaging. bottom, the distribution of the tuning width measured from the calcium or glutamate imaging. No significant difference was found in any comparison. **f,** The probability to find a responsive spine in iGluSnFR3 is three times more than in GCaMP8s imaging (n/N = 913/7, Wilcoxon rank-sum test, *P < 0.001). **g,** After alignment of the soma across multiple cells (N = 21), the centroid of the distribution of spines receiving the inputs from the information likely supplied by different mechanosensitive end organs reveals the similar rostral-caudal bias of the end organ types. Number of experiments and statistics in Extended Data Table 1-2.

**Extended Data Figure 8.**
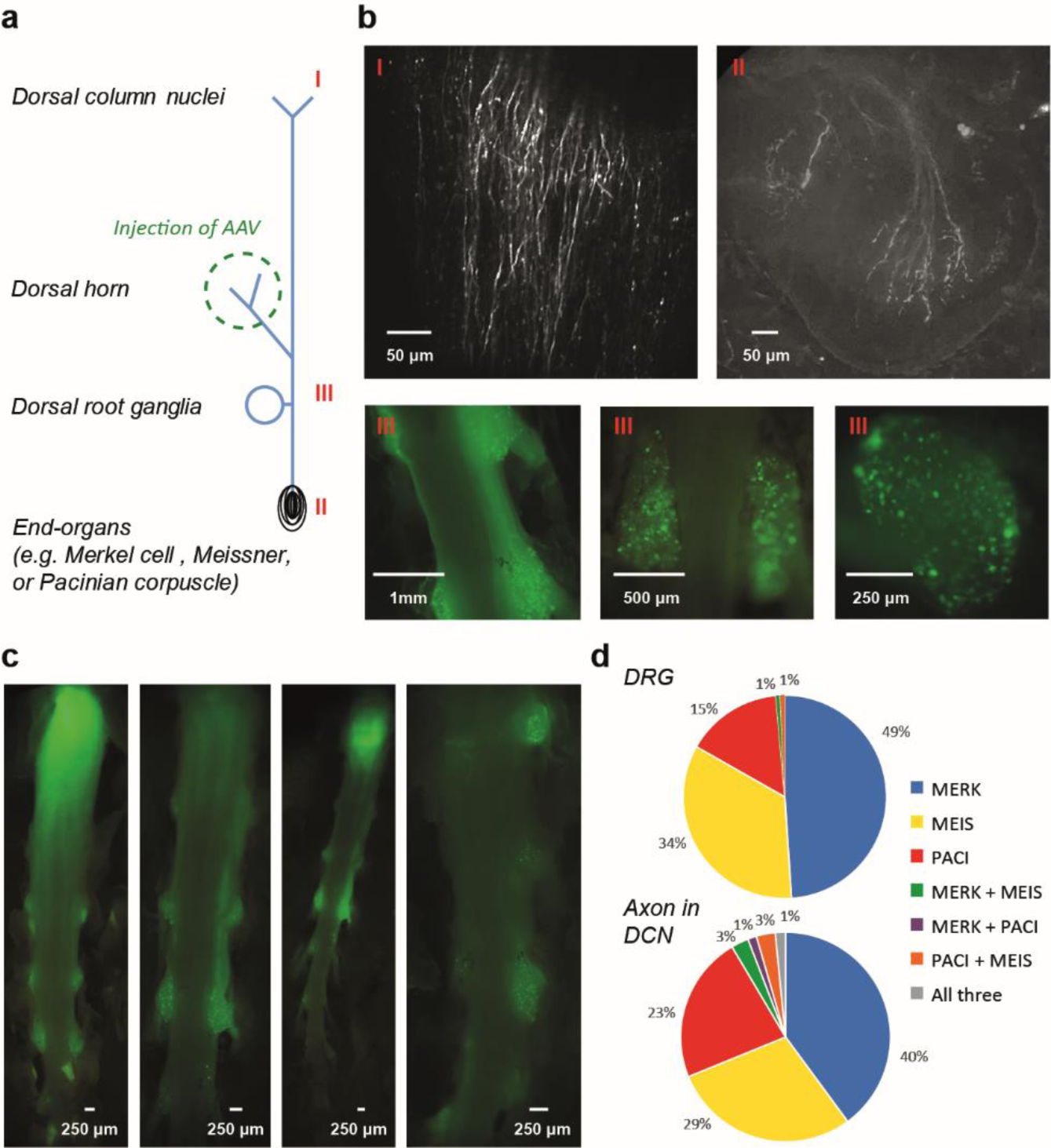
Details about tagging first-order somatosensory neurons and axonal bouton imaging. **a,** Expression of GCaMP6s in the first-order somatosensory neurons in DRG was achieved by viral injection at the dorsal horn of lumbar enlargement. **b,** The two photon imaging of the brainstem (top left) and a whole-mount skin sample from the toe tip (top right) showed the two extreme terminals of the first-order somatosensory neurons. Bottom, three example fields of view at the DRG level, showing the labelling of the cell bodies. **c,** Four example images of dissected spinal cord revealed the heterogeneity of the viral injection and expression. **d,** Top, the percentage of different groups of DRG neurons defined by three mechanosensitive end organs-specific stimulation (n/N = 143/3). Bottom, the percentage of seven classified groups of axonal boutons in DCN defined by three mechanosensitive end organs-specific stimulation (n/N = 446/6). Those multimodal DCN boutons are likely from the indirect pathway through the spinal cord dorsal horn.

**Extended Data Figure 9.**
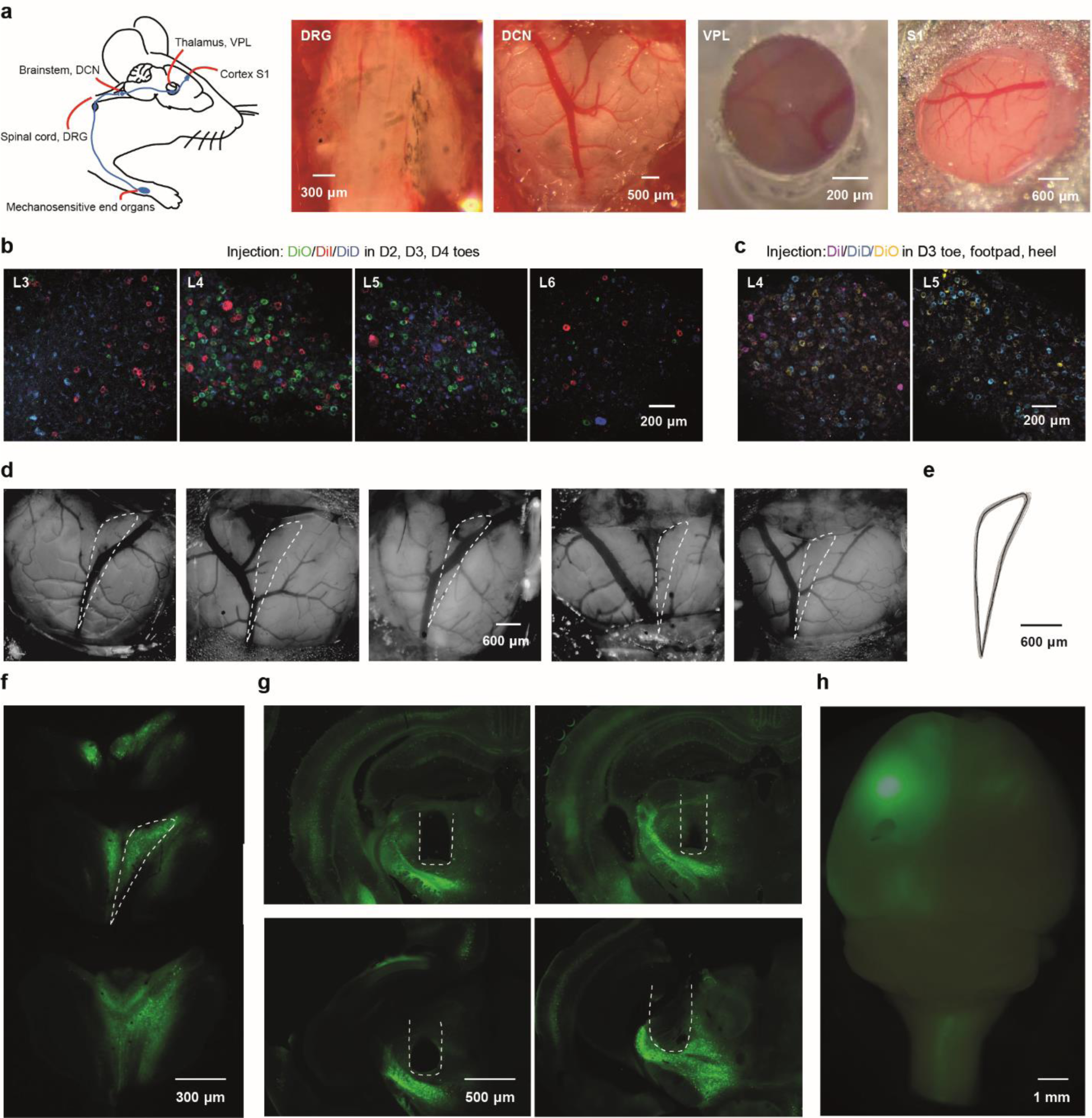
Histology and illustration of imaging in the complete somatosensory pathway for touch. **a,** Left, schematic representation of the ascending pathway of the somatosensory system. Right, the example photos of each location where the imaging windows were installed. **b-c,** The anatomical-defined receptive fields of lumbar DRG cells were mapped by injecting fluorescent lipophilic dye DiI, DiO and DiD into different locations of the hindlimb. **d,** Example photos of five DCN windows. White dashed lines marked the right gracile nucleus. **e,** Overlaying of the gracile nuclei across 24 animals yielded a template for alignment. **f,** Three tangential serial sections (100 μm apart) of the DCN showing the viral expression covered the complete right gracile nucleus. White dashed lines marked the right gracile nucleus. **g,** Four example *post hoc* histological verification of the GRIN lens implantation above the thalamus (VPL) after the viral expression of GCaMP7b. **g,** Example epifluorescence imaging of the brain showing the GCaMP expression at the injection site at S1 and the projections at the contralateral DCN.

**Extended Data Figure 10.**
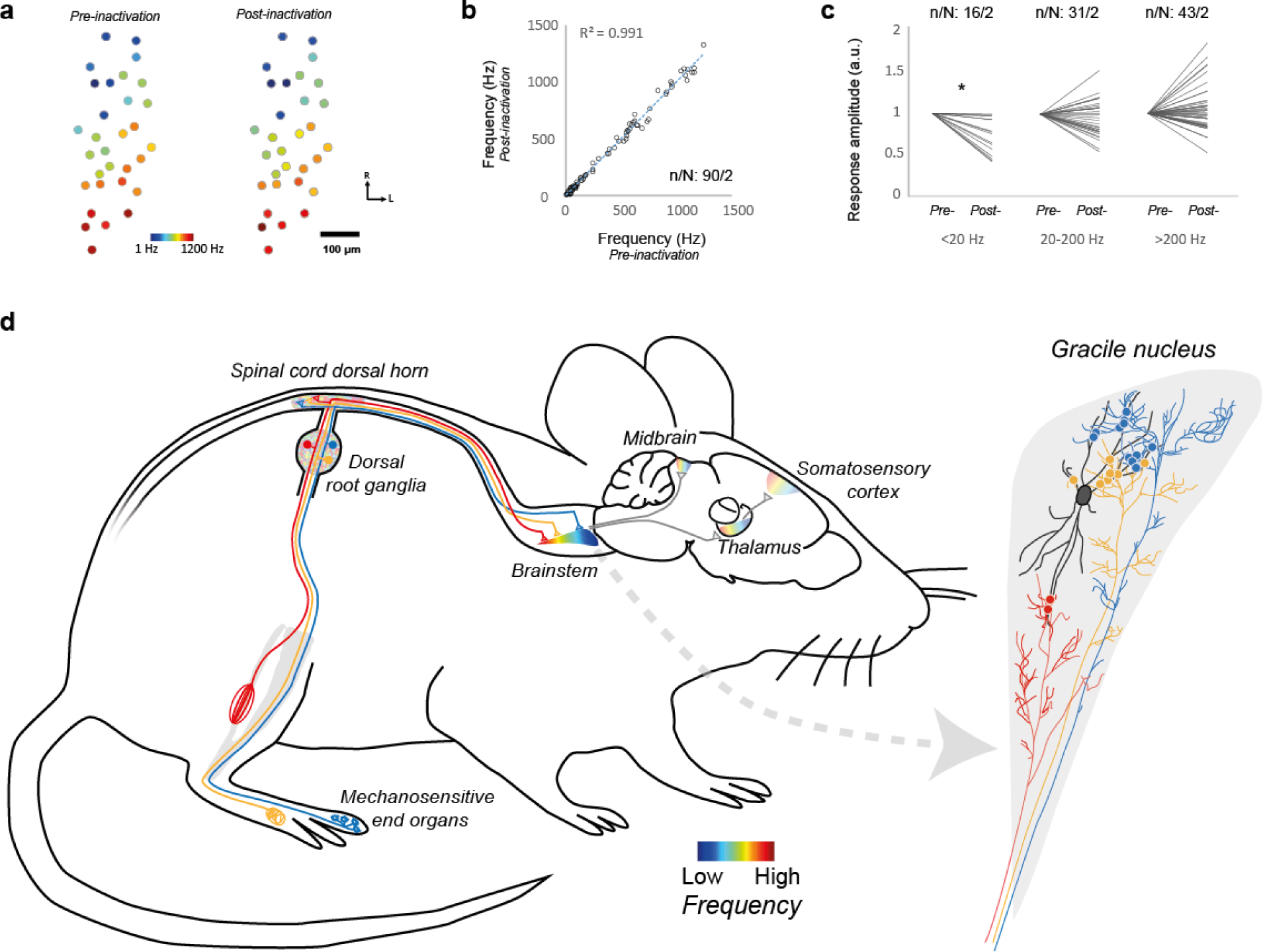
A model for emergence of tonotopic representation in the hindlimb dorsal column–medial lemniscal pathway. **a,** Frequency preference of neurons are colour-coded before (left) and after (right) the application of inhibitors of excitatory synaptic transmission to the spinal cord. **b,** The correlation of the frequency preference of individual neurons measured before and after the inactivation of the synaptic activity within the spinal cord (linear regression, P < 0.001). **c,** The change of the calcium response amplitude of individual neurons in three different groups, categorized by their preferred frequency, before and after the inactivation (paired-sample t-test, *P < 0.001). **d,** Mechanosensitive end-organs and their LTMR afferents - Merkel cells (blue), Meissner corpuscles (yellow) and Pacinian corpuscles (red) - detect a wide frequency spectrum of vibrations (0.1 - 1000 Hz) in the periphery. The end organs are distributed throughout the hindlimb, but each is most densely found in one area: Merkel cells in the toes, Meissner corpuscles in the foot pads and Pacinian corpuscles along the fibula. At the cell bodies of the LTMRs in the dorsal root ganglia no tonotopic organisation is observed. However, at their central projections in the gracile nucleus of the brainstem, fine-scale topographic organisation arises. Furthermore, inputs from the same end-organ group are found projecting onto the gracile nucleus with a rostrocaudal bias. Each colour-coded circle illustrates a synaptic connection. The selective dendritic sampling of the gracile nucleus neurons (black) is the basis for the emergence of a topographic map in the brainstem. Along the ascending pathway, the tonotopic map found in the gracile nucleus is partially preserved in the thalamus and hindlimb somatosensory cortex.

**Extended Data Table 1:** number of experiments.

| **Figure** | **Experiment** | **N (animals)** | **n (units)** |
| --- | --- | --- | --- |
| 1c | Location tuning mapping | 1 | 98 |
| 1d | Frequency tuning mapping | 1 | 98 |
| 1g | Linear somatotopy | 1 | 98 |
| 1h | Linear tonotopy | 1 | 98 |
| 1i | Columnar organization | 7 | 892 |
| 1j | Maps intersection | 7 | 892 |
| 2e | Functional clustering, DRG | 6 | 168 |
| 2f | Functional clustering, DCN | 7 | 892 |
| 2g | Functional clustering, VPL | 5 | 87 |
| 2h | Functional clustering, S1 | 4 | 507 |
| 2i | Freq&location joint representation, DRG | 6 | 168 |
| 2j | Freq&location joint representation, DCN | 2 | 184 |
| 2k | Freq&location joint representation, VPL | 5 | 87 |
| 2l | Freq&location joint representation, S1 | 4 | 507 |
| 3b | Threshold mapping | 10 | 92 |
| 3d | Distribution of end organ type response | 4 | 477 |
| 3e | Percentage of responsive groups | 4 | 477 |
| 4a | Distribution of feature preference, axon | 7 | 913 |
| 4b | Functional clustering, axon | 7 | 913 |
| 4c | Distribution of feature preference, spine | 6 | 410 |
| 4d | Functional clustering, spine | 6 | 410 |
| 4f | Functional specificity of connections | 5 | 7 |
| ED2 | Vibration transmission | 2 | N/A |
| ED3 | Invariance of the receptive field properties | 2 | 81 |
| ED4b | Tuning properties, GCaMP6s | 4 | 424 |
| ED4b | Tuning properties, GCaMP6f | 3 | 164 |
| ED4b | Tuning properties, GCaMP7b | 2 | 159 |
| ED4b | Tuning properties, GCaMP8s | 6 | 504 |
| ED4c | Tuning properties of DRG, GCaMP6s | 6 | 212 |
| ED4c | Tuning properties, electrophysiology | 5 | 84 |
| ED5a | Distribution of Pacinian corpuscle | 5 | 243 |
| ED7b | Percentage of responsive groups, spines | 7 | 358 |
| ED7b | Percentage of responsive groups, cells | 11 | 21 |
| ED7d | Functional clustering, along dendrites | 5 | 205 |
| ED7e | Tuning properties, GCaMP8s | 3 | 102 |
| ED7e | Tuning properties, iGluSnFR3 | 4 | 246 |
| ED7g | Centroid of responsive groups | 7 | 223 |
| ED8d | Percentage of responsive groups, DRG | 3 | 143 |
| ED8d | Percentage of responsive groups, axon | 6 | 446 |
| ED9e | Alignment of DCN | 24 | N/A |

**Extended Data Table 2:**
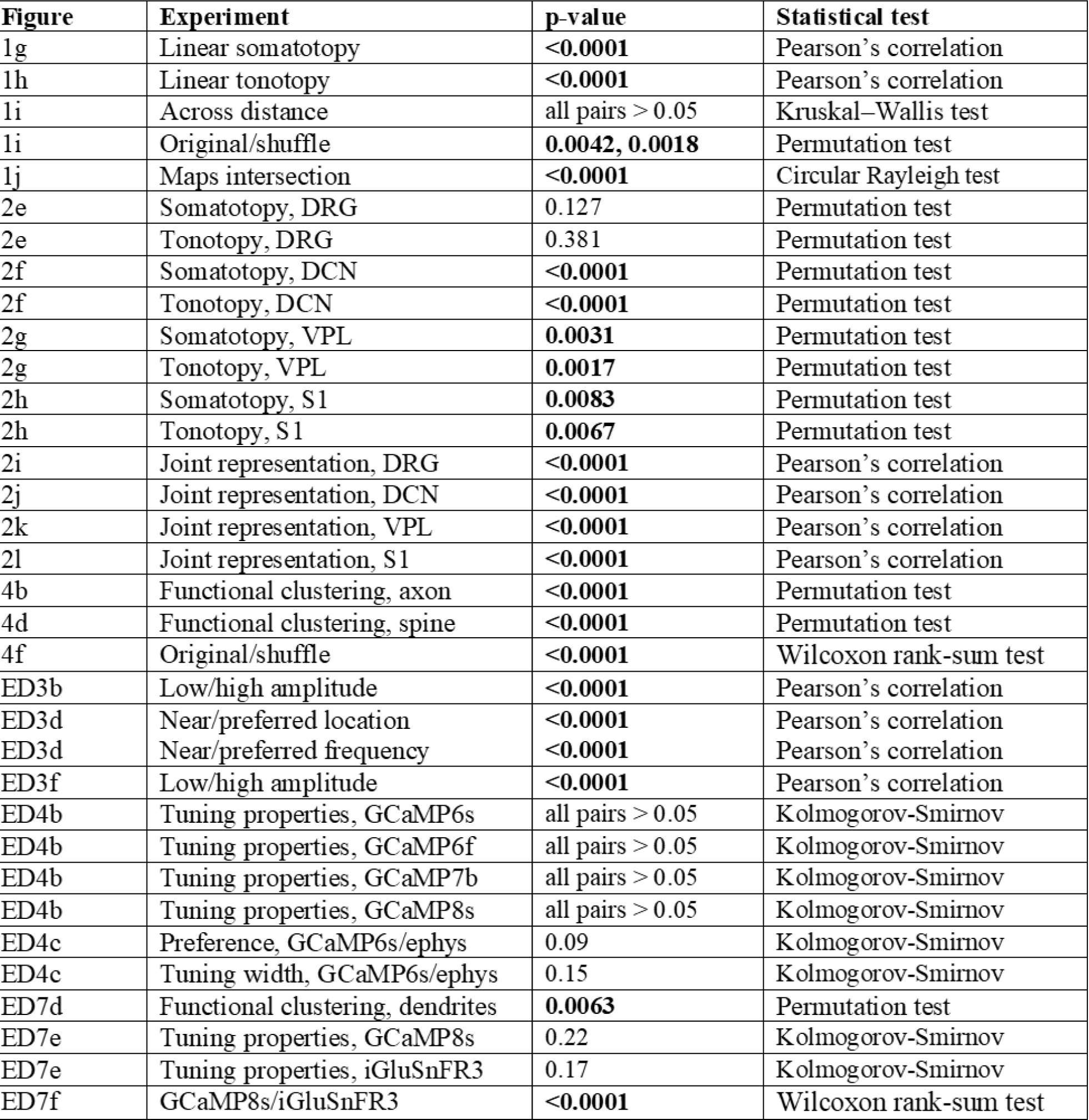
statistics.

| **Figure** | **Experiment** | **p-value** | **Statistical test** |
| --- | --- | --- | --- |
| 1g | Linear somatotopy | **<0.0001** | Pearson's correlation |
| 1h | Linear tonotopy | **<0.0001** | Pearson's correlation |
| 1i | Across distance | all pairs > 0.05 | Kruskal–Wallis test |
| 1i | Original/shuffle | **0.0042, 0.0018** | Permutation test |
| 1j | Maps intersection | **<0.0001** | Circular Rayleigh test |
| 2e | Somatotopy, DRG | 0.127 | Permutation test |
| 2e | Tonotopy, DRG | 0.381 | Permutation test |
| 2f | Somatotopy, DCN | **<0.0001** | Permutation test |
| 2f | Tonotopy, DCN | **<0.0001** | Permutation test |
| 2g | Somatotopy, VPL | **0.0031** | Permutation test |
| 2g | Tonotopy, VPL | **0.0017** | Permutation test |
| 2h | Somatotopy, S1 | **0.0083** | Permutation test |
| 2h | Tonotopy, S1 | **0.0067** | Permutation test |
| 2i | Joint representation, DRG | **<0.0001** | Pearson's correlation |
| 2j | Joint representation, DCN | **<0.0001** | Pearson's correlation |
| 2k | Joint representation, VPL | **<0.0001** | Pearson's correlation |
| 2l | Joint representation, S1 | **<0.0001** | Pearson's correlation |
| 4b | Functional clustering, axon | **<0.0001** | Permutation test |
| 4d | Functional clustering, spine | **<0.0001** | Permutation test |
| 4f | Original/shuffle | **<0.0001** | Wilcoxon rank-sum test |
| ED3b | Low/high amplitude | **<0.0001** | Pearson's correlation |
| ED3d | Near/preferred location | **<0.0001** | Pearson's correlation |
| ED3d | Near/preferred frequency | **<0.0001** | Pearson's correlation |
| ED3f | Low/high amplitude | **<0.0001** | Pearson's correlation |
| ED4b | Tuning properties, GCaMP6s | all pairs > 0.05 | Kolmogorov-Smirnov |
| ED4b | Tuning properties, GCaMP6f | all pairs > 0.05 | Kolmogorov-Smirnov |
| ED4b | Tuning properties, GCaMP7b | all pairs > 0.05 | Kolmogorov-Smirnov |
| ED4b | Tuning properties, GCaMP8s | all pairs > 0.05 | Kolmogorov-Smirnov |
| ED4c | Preference, GCaMP6s/ephys | 0.09 | Kolmogorov-Smirnov |
| ED4c | Tuning width, GCaMP6s/ephys | 0.15 | Kolmogorov-Smirnov |
| ED7d | Functional clustering, dendrites | **0.0063** | Permutation test |
| ED7e | Tuning properties, GCaMP8s | 0.22 | Kolmogorov-Smirnov |
| ED7e | Tuning properties, iGluSnFR3 | 0.17 | Kolmogorov-Smirnov |
| ED7f | GCaMP8s/iGluSnFR3 | **<0.0001** | Wilcoxon rank-sum test |

## Methods

### Animals

Imaging and electrophysiology experiments were conducted with C57BL/6 (Charles River Laboratory) mice. Hindpaw anatomical localization of Pacinian corpuscle experiments were conducted in double-transgenic mice generated by mating homozygote Ai14 males carrying a floxed tdTomato fusion gene inserted in the *Gt(ROSA)26Sor* locus in a C57BL/6 strain ^38^ (Jackson Laboratory; stock no. 007914) with heterozygote ER81/Etv1-CreER females expressing the CreER^T2^ fusion protein from the *ER81/Etv1* promoter elements ^39^ (Jackson Laboratory; stock no. 013048). As such, Cre-mediated recombination resulted in an expression of the floxed tdTomato sequence in the ER81/Etv1-expressing cells of the offspring. ER81/Etv1 is expressed in the inner core region of Pacinian corpuscles ^40^. To induce CreER-based recombination, double-transgenic adult offspring were administered one daily dose of a 100 µL tamoxifen (T5648, Sigma-Aldrich) solution (20 mg/mL) dissolved in corn oil (C8267, Sigma-Aldrich) for five consecutive days by peritoneal injection. Mice were housed in an animal facility, maintained on a 12:12 light/dark cycle, and the experiments were performed during the light phase of the cycle. The animals did not undergo any previous surgery, drug administration or experiments and were housed in groups of maximum 5 animals per cage. All procedures complied with and were approved by the Institutional Animal Care and Use Committee of the University of Geneva and Geneva veterinary offices.

### Surgery

15 to 30 week old mice were surgically prepared for terminal two-photon Ca2+ imaging. Surgeries were conducted under isoflurane anaesthesia (1.5 to 2%) and additional analgesic (0.1 mg/kg buprenorphine intramuscular (i.m.)), local anaesthetic (75 µL 1% lidocaine subcutaneous (s.c.) under the location for incision) and anti-inflammatory drugs (2.5 mg/kg dexamethasone i.m. and 5 mg/kg carprofen s.c.) were administered. Mice were fixed on a bite bar with a snout clamp and rested on top of a heating pad. For viral expression at the brainstem, thalamus and cortex, the scalp was cut over the midline between the ears and eyes. After removing the dura, a bulk viral injection (400 to 800 nL) was administered with pulled and bevelled glass pipettes (≈15 µm tip diameter) at an approximate rate of 20 nL/min into either, the gracile nucleus right next to the obex, the ventral posterolateral nucleus (VPL), nucleus of the thalamus (2 mm lateral and 2 mm posterior to bregma; 3.5 mm below the dura), or the hindlimb representation of the primary somatosensory cortex (2 mm lateral and 0 mm to anterior to bregma; 350 µm below the dura). For a dense expression among the neural population, the injections consisted of AAV1.Syn.GCaMP6f.WPRE.SV40 (University of Pennsylvania, CS0939, titer 6.93 x 10^13^), AAV1.Syn.GCaMP6s.WPRE.SV40 (University of Pennsylvania, CS0205, titer 3.22 x 10^13^), or AAVrg.Syn.jGCaMP7b.WPRE (Addgene, lot. no. v63074, titer 1 x x 10^13^). For sparse expression of GCaMP in the brainstem neurons, AAV1.hSyn.Cre.WPRE.hGH (University of Pennsylvania, CS0938, titer 5.05 x 10^13^) was diluted (1:100,000) in phosphate-buffered saline (Sigma) and mixed with AAV1.CAG.Flex.GCaMP6s.WPRE.SV40 (Addgene, lot. no. v24793, titer 1.9 x 10^13^), AAV1.Syn.FLEX.jGCaMP8s.WPRE (Addgene, lot. no. v111208, titer 2.3 x 10^13^), AAV1.hSyn.FLEX.iGluSnFR3.v857.PDGFR (University of Zurich, titer 7.3 x 10^13^), or mixed with AAV5.hSynap.FLEX.SF-iGluSnFR.A184S (Addgene, titer 1.3 x 10^13^). For GCaMP expression in the DRG neurons and their axonal terminals within the DCN, AAV5.CAG.GCaMP6s.WPRE.SV40 (Addgene, lot. no. v20751, titer 3.2 x x 10^13^), or AAV1.CAG.GCaMP6s.WPRE.SV40 (University of Pennsylvania, V3096TI-3, titer 2.33 x 10^13^) was directly injected (1000 to 3000 nL) into the lumbar enlargement in between T13 and L4 vertebrates. For visualising the process of injection, 0.1 μL of 2% FastGreen was added into 5 μL of the virus. After the end of each injection, the pipette was left in place for at least another 10 min before being retracted. After 3-8 weeks of expression and recovery, experiments could be conducted.

All imaging and electrophysiology experiments were surgically prepared under terminal anaesthesia by inhalation of isoflurane (∼2%) and the body temperature was maintained near 37 °C. For analgesia, buprenorphine (0.1mg/kg SC) was provided 15-20 min before the procedure, except for thalamus and cortex, where chronic preparation was employed. An additional dose of buprenorphine was given if the procedure exceeded 4 hours. The whole procedure was limited to 8 hours. For imaging of brain structures, a custom-made titanium head bar was fixed on the skull with a cyanoacrylate adhesive (ergo 5011, IBZ Industrie) and dental cement to allow subsequent head fixation.

Imaging of dorsal root ganglia (DRG) was done at least four weeks after GCaMP expression. The surgical procedure for vertebral window implantation was adapted from a previous study ^41^. In short, after hemostasis, DRG was covered with a thin layer of Kwik-Sil silicone elastomer (World Precision Instruments) and sealed with a 2-mm diameter cover glass, and the cover glass was attached to the vertebral mount by cyanoacrylate glue and dental acrylic.

For brainstem imaging, the skin and cervical paraspinal muscle over the neck were cut and pulled to the side. A custom made titanium head bar was fixed on the skull with a cyanoacrylate adhesive (ergo 5011, IBZ Industrie) and dental cement. The head was fixed at approximately 45° downward. After the removal of dura and the skull above the posterior part of the cerebellum, Kwik-Sil was applied to the surface of the brainstem before a hand-cut wedge-shaped glass coverslip (150 µm thickness) was placed over the brainstem. The glass window over the gracile nucleus was first stabilised by the pressure of a micromanipulator. The micromanipulator was removed after the cyanoacrylate glue and dental acrylic hardened.

For thalamic and cortical imaging, survival surgery for window implantation was performed either right after viral injection or after 2-4 weeks of recovery period and viral expression. A craniotomy was performed over the target structures. For the cortical imaging, two hand-cut glass coverslips (150 µm thickness) matched to the shape of the craniotomy were glued together with optical adhesive (NOA 61, Norland Products). The lower glass was placed on top of the cortex, and the upper glass, which was larger than the craniotomy, was glued to the skull with cyanoacrylate adhesive and secured with dental cement. For thalamic imaging, a gradient-index (GRIN) lens (Inscopix) was used instead of a glass coverslip. The correspondence between the injection sites and the sensory hindpaw representation was confirmed during *in vivo* sensory mapping.

### Two-photon Ca^2+^ imaging

Imaging was performed with a custom-built two-photon microscope (MIMMS, www.openwiki.janelia.org) controlled by Scanimage 5.1 (Vidrio Technologies) using a 16× 0.8 NA objective (Nikon) and with excitation wavelength set to ∼920 nm (Ultra II, tunable Ti:Sapphire laser, Coherent). 512 by 512 pixel images were acquired at 29.72 Hz using bidirectional scanning with a resonant scanner system (Thorlabs). The power was modulated with a Pockels cell (350-80-LA-02, Conoptics) and calibrated with a photodiode (Thorlabs). The primary mirror for imaging was a custom polychroic (Chroma, zt470/561/nir-trans) that could transmit infrared light, while reflecting green light. Before detection, the remaining infrared light was filtered with a coloured glass band pass filter (BG39, Chroma). Images were acquired using photomultiplier tubes (H11706P-40 SEL, Hamamatsu) and written in 16-bit format to disk.

For dendritic imaging, individual neurons in DCN were selected for imaging based on several criteria: visible dendritic spines, nuclear exclusion, responses to tactile stimulation, and a lack of large blood vessels obscuring the dendritic field ^42^. Images of dendritic segments were acquired at 29.82 Hz (resolution: 8.42–12.2 pixels/μm) and z-stacks of individual cells were acquired prior to dendritic imaging by averaging 50 frames per plane using 1–2 μm z-steps. Multiple dendrites across multiple depths were imaged on individual cells. This imaging method only allows visualisation of a fraction of spines on dendrites in horizontal planes. Similar logic applied to imaging of axonal boutons and genetically encoded glutamate indicators (128 by 128 pixel images, at 100.23 Hz) in both dendrites and axons. Throughout the experiment, dendrites and axons were carefully monitored for indications of photodamage.

Raw calcium images were processed using Suite2P, a publicly available two-photon calcium imaging analysis pipeline ^43^. First, images were registered based on the cross-correlation of the averaged images to account for brain motion. Then, regions of interest (ROI) were established by clustering neighbouring pixels with similar time courses. Next, manual selection of ROI was performed to eliminate low-quality or non-targeted regions of interest. Each field of view was aligned to the photograph of the dorsal surface of DCN using an affine transform, which was applied in ImageJ (NIH). In some cases, DCN fields of view across multiple animals or from both hemispheres were aligned to a template of the averaged DCN images from the right side.

### Electrophysiology

For all electrophysiology experiments, the mice were anaesthetised by isoflurane inhalation (2%) and body temperature was maintained near 37 °C with a heating mat. For analgesia, buprenorphine (0.1mg/kg SC) was provided 15-20 min before the procedure.

To assess the mechanosensitive properties of the afferents of Merkel cells, Meissner corpuscles and Pacinian corpuscles, first, a skin opening was performed along the hamstring muscle. Then, the sciatic nerve was carefully isolated using ophthalmic scissors and tweezers. The pia mater spinalis and dura mater around the nerve were removed, and finally, the nerve bundle was separated into single fibres (15-20 μm in diameter) and placed on two Ag/AgCl wire electrodes in the recording pool filled with mineral oil. A third ground Ag/AgCl electrode was placed in the muscle next to the recording chamber.

To characterise the response properties of sensory afferents at the peripheral level, we recorded stimulus-evoked activity of afferents *in vivo* using hook electrodes in the tibial nerve. Vibratory stimuli were applied to different locations of the hindlimb using a calibrated piezo stimulator (Physik Instrumente P-841, E509 controller, E504 amplifier), so that the neural responses to all combinations of locations, frequencies and amplitudes could be characterised. To determine the mechanical sensitive threshold and the frequency tuning of each afferent, a series of sinusoidal mechanical stimuli (duration 20 s for frequencies above 100 Hz and correspondingly increasing for lower frequency; linearly increasing amplitude) were applied to the hand-mapped receptive field of individual afferents with the piezo actuator. In order to study pure vibration induced response, the probe was placed on the stimulation location for at least 10 seconds and the stimulation for threshold mapping ramping up slowly. Thus, the adaptation plays a role.

The signal was amplified and filtered (>10 Hz and <10 kHz) and acquired at 30 kHz (PXIe-1073, National Instruments) using WaveSurfer (https://wavesurfer.janelia.org/) Matlab (Mathworks) routines. Trial start, stimulus onset triggers, and details of stimulus were saved in parallel on separate channels and used for *post-hoc* alignment of recorded spikes. After the recording, animals were euthanized by overdose with isoflurane (5%) followed by cervical dislocation and bleeding.

### Pharmacology

For silencing the indirect dorsal column pathway, animals were prepared for DCN two-photon calcium imaging as described above. The surgery at the spinal cord area was similar to the vertebral window implantation for the imaging of dorsal root ganglia. After a laminectomy over L3–L6 spinal segments, Kwik-Sil silicone elastomer (World Precision Instruments) was used to create a pool and confine drugs to the spinal cord. The dura was removed from the spinal cord using fine forceps. Drug delivery followed the previous established protocol ^9^. The non-competitive NMDA receptor antagonist MK-801 (10 mM, 10 µl, Abcam; dissolved in 90% H2O and 10% DMSO) was applied to the surface of the spinal cord and allowed to enter the cord for 3–5 min. The surface of the cord was then irrigated with saline. NBQX (10 mM, 20 µl, Abcam, dissolved in H2O) was then applied to the surface of the cord. After 5 min, the cord was covered with gelfoam, which was allowed to absorb the NBQX and remained in place for the duration of the experiment, occasionally re-wet with saline. After the calcium imaging, and animals were sacrificed within 2– 3 h following drug application.

### Vibrotactile stimuli

The vibrotactile stimuli were generated by a bimorph piezoelectric multilayer bender actuator with either a pre-attached holder (Thorlabs, PB4NB2S) for low frequency stimulations (below 100 Hz) or a piezoelectric stack actuator (P-841.3, Physik Instrumente) equipped with a strain gauge feedback sensor for high frequency stimulations (above 100 Hz). A hand-cut blunt plastic cone was mounted on the actuator. The actuator and sensor controllers (E-662, E-618.1 and E-509.S1, Physik Instrumente) operated in closed loop, thereby counteracting the contacting force on the skin of mice (below 50 mN). The forces of our vibrotactile stimuli measured at the static states of 3µm, 30 µm and 300 µm, were less than 10mN, 10-30 mN, and around 50mN, respectively. Pure sinusoids (250 or 500 ms duration) of a wide range of frequencies (0.1 to 3000 Hz) calibrated to produce a desired displacement amplitude (0.01 to 1000 µm) were sampled at 10 kHz (USB-6353, National Instruments) and fed to the actuator controller. The sensor measurements were continuously acquired and recalibration of motor commands was regularly performed for the stimuli to remain highly consistent, by comparing the ground truth data acquired optically with a laser doppler vibrometer system (Polytec, OFV-5000). The spectrum of the acquired sensor measurements was analysed to ensure the integrity of their frequency content (**Extended Data Fig. 1**).

To map the location tuning curve of each neuron, a 100 Hz vibration at an amplitude of 400 µm (for 500 ms) was applied at seven locations of the hindlimb (**Fig. 1a**). These frequency and amplitude parameters were selected as they drive the LTMRs that innervate all three major mechanosensitive end organs. To measure the frequency tuning curve of each neuron, three sets of stimulations were applied at three locations of the hindlimb. The three locations were equivalent to the second, fourth and sixth positions used in the location preference mapping (**Fig. 1a**). The three sets of stimulations were done at the amplitudes of 300 µm (testable bandwidth: 0.1-100 Hz), 30 µm (testable bandwidth: 30–600 Hz) and 3 µm (testable bandwidth: 100–3000 Hz), which correspond to the sensitive ranges of the LTMRs innervating each Merkel cells, Meissner corpuscles and Pacinian corpuscles, respectively. At each amplitude, seven to eleven different frequencies were chosen incrementally and repeated for four to eight trials. Of the three vibration amplitudes presented, the one that evoked the greatest neural response was identified for each neuron. The stimulus set of that given amplitude was then used to determine the frequency tuning curve for that neuron for further analysis. To characterise the input type, the end organ-specific stimuli (Merkel cell, 1 Hz, 300 μm; Meissner, 30 Hz, 100 μm; Pacinian, 600 Hz, 3μm) were applied at the all seven locations of the hindlimb (**Fig. 1a**).

### Vibration conductance

Vibrations transmitted within the hindlimb of the mice were measured using a scanning laser Doppler vibrometer (Polytec, OFV-5000) at distances between 5 and 20 cm, either on the skin of the anaesthetised mice or on the fibula of a freshly dissected hindlimb *post mortem* (**Extended Data Fig. 2**). Piezoelectric stack actuators were either attached to the toes or the heel. The trigger signal was generated internally with a customised code in Matlab. Both vibrometer outputs and the trigger signal were recorded simultaneously on a laptop computer. For the seven measurement points along the hindlimb, the position coordinates and the displacement data of individual trials were extracted. Since the vibrometer output voltage was proportional to the arbitrary displacement, these values can be translated back to actual displacement for the analysis of scanning trials.

### Histology

Upon completion of terminal experiments, isoflurane was raised to 5% and 0.2 ml pentobarbital was delivered through an intraperitoneal injection. The animal was transcardially perfused with 20 ml of 0.9% NaCl and then 50 ml of 4% paraformaldehyde in 0.1 M PB. The brain or body parts were removed and placed in 4% PFA overnight. A vibratome (Leica VT1200S) was used to cut tangential or coronal sections (100 µm thick). Tangential sections were cut parallel to the in vivo imaging window surface to aid in *post hoc* identification of previously imaged structures *in vivo*. Slices were washed three times using 1X PBS, then mounted to a slide using SlowFade Gold (Thermofisher Scientific) or Mounting Medium with DAPI (Ancam,104139) for imaging using a fluorescence microscope (Olympus BX53) or confocal microscopy (ZEISS LSM 800).

### Epifluorescence and confocal imaging

For *in situ* imaging, skin, muscles and fat were resected to expose the area of interest in the limbs or spinal cord. This was done under a fluorescent microscope (Leica M165FC). The expression pattern of GCaMP or tdTomato was imaged by illuminating the dissected tissue with a blue or green LED light, with emission channels optimised for each fluorophore (GRF, or RFP). In some cases, the *in situ* imaging of Pacinian corpuscles in the toes or hindlimb was also done using two-photon microscopy to achieve higher resolution. Furthermore, once the fibula was exposed by resection of skin and muscles, the interosseous nerve and Pacinian corpuscles located on the bone could be excised and subsequently imaged with confocal or two-photon microscopes.

Confocal imaging was done using a ZEISS LSM 800 confocal microscope. Images were acquired using 405, 488, 561, and 633 nm laser lines with emission channels optimised for each fluorophore (DiI, DiD, DiO, GRF, and tdTomato) while minimising cross-talk between channels. Images were acquired at 512×512 with sampling resolution ranging from 0.15 to 1 μm per pixel. Z-stacks were acquired using 0.5 to 5 μm steps and the confocal pinhole was set to 1 AU. Colocalization was performed manually. The imaging data was analysed using ImageJ (NIH).

### Data analysis

#### Significant responses

ROI drawing for the targeted neuronal structures was performed in ImageJ (NIH). For somas, boutons and spines, ROIs were circular or drawn manually. For the dendrites, polygonal ROIs were drawn spanning the extent of a short, contiguous dendritic segment. Spine distances on dendritic segments were measured by reconstructing dendritic arbours using Simple Neurite Tracer. Fluorescence time-courses were computed as the mean of all pixels within the ROI at each time point, then extracted using Miji ^44^, and finally synchronised with the stimulation. Evoked responses were computed as changes in fluorescence relative to baseline fluorescence. Dendritic-spine calcium signals were sometimes contaminated by regenerative dendritic events. We used a subtraction procedure to isolate spine signals: (1) performing a robust fit (MATLAB) of the spine signal against the dendritic signal for stimulus-evoked data and (2) subtracting a scaled version of the dendritic signal, where the scaling factor equals the slope from the robust fit ^16^. Following subtraction, dendritic spines correlated with dendritic signals (r > 0.50) were excluded from analysis.

Location and frequency selectivity for individual neurons, axonal boutons and dendritic spines was quantified by a previously reported method ^19^. To computing tuning properties, the fluorescence signal was calculated as Δf/f=(f–f0)/f0, where f0 is the baseline fluorescence signal averaged over the 1 s period immediately before the start of the vibration stimulation and f is the fluorescence signal averaged over the first 1.5 s period after the start of the stimulation. Tuning curves were obtained by calculating the mean fluorescence signal (Δf/f) for each stimulation, and then fitting a polynomial curve to the resulting data.

#### Tuning curve fitting

Neuronal structures were considered to be responsive if the maximum stimulus-related fluorescence response (Δf/f) to any stimulus was greater than 5% on average, and also greater than 2 standard deviations (SD) above the mean baseline fluorescence. In addition, we required that neurons responded at least 2 SD above baseline on at least 20% of the trials tested. Neuronal structures were considered to be location and frequency selective if they were responsive and also met the following criteria: (1) well fit by the polynomial function (r > 0.7, P < 0.05), and (2) tuning index (*TI*) > 0.2

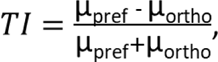

where μ_pref_ denotes the mean response to the preferred stimulation and μ_ortho_ is the mean response to the least preferred stimulation, defined by the curve fitting.

To characterise a neuron’s tuning to vibration frequency and location, the normalised mean responses were fit to frequency by the polynomial curve fitting function (Matlab) using the method of non-linear least squares, with the limited degree at 6. The tuning preference of individual neurons was defined by the peak of the curve and the tuning width was defined by the half-width at the half-maximum of the curve.

#### Spatial analysis

To summarise the spatial arrangement of different features, the centroid of the distribution of a population of neurons with a certain responsive profile toward the stimulation was depicted. The vector and the axis of somatotopy was defined by the centroid of neurons responsive to stimulation at the two most distant sites: the toe and thigh. It was used to demonstrate the linear organisation for both the location and frequency along this arbitrary somatotopic axis in DCN. The vector of tonotopy was defined by the centroid of neurons responsive to low frequency stimulation (10 Hz) and to high frequency stimulation (1000 Hz). For the analysis related to the spatial arrangement of the peripheral input, the vector of end organ-specific inputs was defined by the centroid of neurons responsive to Merkel cells-specific stimulus and to Pacinian corpuscles-specific stimulus. The same logic was applied to the analysis of spines within the dendritic fields.

#### Intersection of somatotopic and tonotopic maps

To characterise how the two maps of location and frequency preference are aligned with respect to each other, we first used data obtained from the cellular imaging experiments to characterise how somatotopic and tonotopic maps intersect. The griddata function (Matlab) was used to interpolate 2-D scattered data of neuronal tuning. In some cases, smoothing the raw data with a two-dimensional Gaussian (σ = 25 μm) was performed before computing contours and gradients. To quantify the preferred angle of intersection, we first took the difference in the vector angle of the two map gradients at each pixel. The preferred intersection angle of the gradients was then computed as a function of the intersection histogram ^45^:

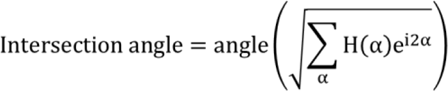

where H(α) is the histogram value as a function of the intersection angle. Note that α is a direction (0° to 360°). A Rayleigh test confirmed that H(α) was significantly different from uniform for all regions.

#### Functional clustering analysis

To characterise the statistical significance of functional clustering among different neuronal structures (from somas to dendritic spines and axonal boutons), we compared the relationship between the pairwise distance of two ROIs and the difference in their preference (location or frequency). In the shuffled data, preferred features were randomly permuted within the brain regions or individual cells. This procedure was repeated 10,000 times to derive the P values for each binned distance. Only the fields of view with at least four feature selective neuronal structures were included in the analysis. Functional clustering was defined as the distance when the experimentally derived Δlocation or Δfreqeuncy was significantly less than that of the shuffled results.

#### Columnar analysis

For analysing the functional organisation in the vertical dimension of the DCN, we applied the same functional clustering analysis to the neurons within the column with a diameter of 30 µm and only included the difference of their difference of vertical distance estimated from the two-photon z-stacks. Whenever we probed the feature selectivity of a column, we always first aligned and collapsed multiple cortical depths (at least three) from the same two-photon field of view into a 2D field of view for straightforward visualisation.

#### Feature extraction and grouping

Automated unsupervised clustering for nerve fibre types was performed on the response profiles to vibration thresholds of different frequencies from 10 to 300 Hz at the receptive field centres. The threshold curves were mapped into a two-dimensional space with t-distributed stochastic neighbour embedding (t-SNE) using the t-SNE Matlab toolbox (https://lvdmaaten.github.io/tsne). The initial dimensionality reduction performed by t-SNE was set to 5 dimensions and the t-SNE perplexity parameter was set to 30. Four different distance metrics (Chebychev, Euclidian, cosine and Mahalanobis) were tested, and cosine distance was chosen because this metric resulted in the best cluster quality. The mapped points formed identifiable clusters, each corresponding to the sensitivities of different vibration frequencies (Extended Data Fig. 6f).

#### Dendritic model

To test the hypothesis of random input sampling of DCN neurons underlying the emergence of the tonotopic map, cellular input–output transformations were computed either based on the actual measured synaptic tuning properties or a random sampling from all the axonal boutons located within the dendritic field, which were pooled across different animals. We took the arithmetic mean of dendritic spine responses of all active, frequency-tuned dendritic spines on a cell, then normalised from 0 to 1. We did the same for the individual somas. We then fit frequency tuning curves to these summed responses. We set the preferred frequency for summed spine input to the somatic frequency preference to examine predictability of frequency selectivity between spines and soma. We interpolated the tuning curves and then again normalised the curve from 0 to 1. Finally, we regressed somatic output against summed synaptic input, yielding the linear correlation coefficient r. The fraction of unexplained variance, or nonlinearity, was defined as 1 – r^2^. We performed this same analysis on random sampled tuning curves from the axonal boutons located within the dendritic field. This computation was repeated 1000 times for randomly sampled tuning curves with the same number of the actual dendritic spines measured from the same neuron.

### Statistics

No statistical methods were used to predetermine sample size and all experimental animals were included in the analysis. The normality assumption was tested with the Kolmogorov-Smirnov test. Non-parametric tests were used when the normality assumption was not met. We used a two sided non-parametric Wilcoxon rank-sum test to compare two groups and the Kruskal–Wallis test to compare multiple groups with *post-hoc* tests using Dunn’s test, without assumptions of normality or equal variances. A permutation test was applied to compare the experimentally derived data to shuffled data in the analysis for functional clustering. Pearson’s correlation coefficient was applied to visuotopic and spatial frequency. The Rayleigh test was used to test the uniformity of the intersection angle distribution between the two maps. All statistical methods were two-sided. All data analyses were performed with custom written routines in Matlab.

All data analysed and all custom Matlab code used for the analysis in the current study are available from the corresponding author upon reasonable request.

## Acknowledgements

We thank M. Prsa, A. Loutit, G. Galiñanes, A. Novozhilova, A. Taylor and J. Marozeau for advice and comments on the manuscript; G. Cuenu for help with histological techniques; R. Vickery for advice on electrophysiology experiments; C. Dürst for the assist on iGluSnFR3 experiments; M. Prsa for the assist on experimental setup; and G. Cuenu for help with breeding mice. This work was supported by the Swiss National Science Foundation (310030_184829), the European Research Council (OPTOMOT), and the International Foundation for Research in Paraplegia. K.-S.L. is a EMBO Postdoctoral Fellow (ALTF_816-2020).

## Author contributions

K.-S.L. and D.H. conceptualised the study. K.-S.L. and D.H. designed experiments. K.-S.L. ran experiments and analysed data. M.S. helped with GRIN lens implantation. D.T.W. helped with the hindlimb dissections. K.-S.L. and D.H. wrote the manuscript with assistance from D.T.W.

